# Windborne migration amplifies insect-mediated pollination services

**DOI:** 10.1101/2022.02.01.478668

**Authors:** Huiru Jia, Yongqiang Liu, Xiaokang Li, Hui Li, Yunfei Pan, Chaoxing Hu, Xianyong Zhou, Kris A.G. Wyckhuys, Kongming Wu

**Author notes:** Correspondence: Kongming Wu.

## Abstract

Worldwide, hoverflies (Syrphidae: Diptera) provide crucial ecosystem services (ES) such as pollination and biological pest control. Although many hoverfly species exhibit migratory behavior, the spatiotemporal facets of these movement dynamics and their ES implications are poorly understood. In this study, we use long-term (16 yr) trapping records, trajectory analysis and intrinsic (i.e., isotope, genetic, pollen) markers to describe migration patterns of the hoverfly *Episyrphus balteatus* in China. Our work reveals long-range, windborne migration with spring migrants originating in northern China and exhibiting return migration during autumn. Given the substantial night-time dispersal of *E. balteatus*, this species possibly adopts a ‘dual’ migration strategy. The extensive genetic mixing and high genetic diversity of *E. balteatus* populations underscore its adaptive capacity to environmental disturbances e.g., climate change. Pollen markers and molecular gut-analysis further illuminate how *E. balteatus* visits min. 1,012 flowering plant species (39 orders) over space and time. By thus delineating *E. balteatus* trans-regional movements and pollination networks, we advance our understanding of its migration ecology and facilitate the design of targeted strategies to conserve and enhance its ecosystem services.

## Introduction

Migration plays a key role in the evolution and life history of many organisms, with insects being the most abundant, speciose and economically important group of terrestrial migrants (Chapman et al., 2011). Across the globe, billions of insects annually undertake long-range movements. By transporting energy, nutrients and other organisms between distant regions, insect migrants provide a multitude of ecosystem services and disservices (Chapman et al., 2012; Hu et al., 2016; Satterfield et al., 2020). Despite the important socio-ecological consequences of insect migration, research has primarily centered on a handful of large-bodied charismatic species or agricultural pests (e.g., monarch butterflies, locusts; Chapman et al., 2015). For most other taxa, there’s a critical dearth of information.

Hoverflies (Diptera: Syrphidae) are a speciose family of beneficial insects - deemed to be the second most important pollinators after bees (Branquart and Hemptinn, 2000; Rader et al., 2019). The larval stages of many hoverflies are effective predators of homopteran feeders, providing natural biological control across geographies and farming contexts (Tenhumberg, 1995; Tenhumberg and Poehling, 1995). Evidence to-date suggests that hoverfly species are abundant diurnal migrants that deliver ecosystem services in both natural and man-made habitats (e.g., Lack and Lack, 1951; Aubert and Goeldlin de Tiefenau, 1981; Wotton et al., 2019). Moreover, given that (migratory) hoverflies exhibit comparatively stable population numbers and transport pollen over long distances (Wotton et al., 2019), these species potentially can sustain pollination and pest control services in the face of a global insect decline (Sánchez-Bayo and Wyckhuys, 2019; Powney et al., 2019). Yet, though (long-range) dispersal is a central determinant of their survival, hoverfly migration has only been intermittently studied since the 1950s. In order to effectively conserve these organisms and to raise their contribution to (agro-)ecosystem functioning, a more in-depth understanding needs to be gained of hoverfly migration.

In recent years, several new technologies have helped to uncover insects’ seasonal migration patterns and population genetic structure. Stable isotope analysis, molecular genetics, tethered flight mill assays, insect radar and aerial trapping have all yielded insights into the migration behavior of hoverflies (Ouin et al., 2011; Raymond et al., 2013, 2015; Dällenbach et al., 2018; Wotton et al., 2019; Gao et al., 2020). Attempts have equally been made to capture the geographical extent and ecological impacts of hoverfly migration (Wotton et al., 2019). Most of these studies however originate from (a small area within) Europe, while virtually no information is available from other parts of the world. Also, as the nutritional ecology of most species waits to be deciphered, little is known about hoverfly-plant associations and how those are modulated by (long-range) migration dynamics.

In China, approx. 580 hoverfly species have been described. These include the marmalade hoverfly *Episyrphus balteatus* (DeGeer) (Li et al., 2009), a common flower visitor in urban and agricultural settings across the Palearctic realm. Locally, (insect) migration primarily takes place within the East Asia monsoon climatic zone (Drake and Farrow, 1988). Owing to its geographical range, complex topography and diverse agro-ecological conditions, this climatic zone constitutes an exceptional setting to study broad-scale migration dynamics of hoverfly species e.g., as compared to other parts of the globe (Wotton et al., 2019; Menz et al., 2019; Finch and Cook, 2020).

In this study, we employed a suite of novel methodologies to characterize the migration dynamics of *E. balteatus* in Eastern Asia. More specifically, we conducted long-term (16 yr) searchlight trapping to clarify whether *E. balteatus* engages in long-range migration and to describe the ensuing migration patterns. Second, we deployed backward trajectory analysis and stable isotope analysis to infer the *E. balteatus* migration routes and source areas. Third, we employed a population genetics approach to compare the genetic make-up and demographic history of migrant and field-collected individuals throughout China. Fourth, we described *E. balteatus* host plant associations by identifying the pollen grains attached to hoverfly bodies. Lastly, we paired molecular gut content analysis with high-throughput sequencing (HTS) to investigate the spatiotemporal distribution of its (flower) host plants across a broad geographic range. As such, our work characterized *E. balteatus* migration behavior and captured its broader ecological relevance e.g., in terms of flower visitation networks, pollination or natural biological control services.

## Materials and methods

### Light-trapping and field surveys

During 2003-2018, light trapping was conducted every night from April until October at a field station of the Chinese Academy of Agricultural Sciences (CAAS) at Beihuang (BH) island (Shandong, China; 38°24’N; 120°55’E). With a size of approx. 2.5 km^2^, BH is located in the center of China’s Bohai Strait – an important migration pathway for multiple insect species which originate from the agricultural regions of mainland China at min. 40–60 km distances (Feng et al., 2003). High-altitude migrants of various insect species were collected using a vertical-pointing searchlight trap (model DK.Z.J1000B/t, 65.2 cm diam., 70.6 cm high and ∼30° spread angle; Shanghai Yaming Lighting Co., Ltd., Shanghai, China) equipped with a 1 000-W metal halide-lamp (model JLZ1000BT, Shanghai Yaming Lighting Co. Ltd, Shanghai, China) (Feng et al., 2009). This light-trap was mounted on a platform ∼8 m above sea level. Except for events of heavy rain or power outage, the searchlight was switched on at a daily basis from sunset until sunrise. Trapped insects were collected into a 60-mesh nylon net bag – which was positioned under the trap and manually replaced every 2 h throughout the night. Every day, *E. balteatus* specimens were separated, counted and a subset of trapped individuals was individually stored at -20 °C in Eppendorf tubes for further analysis. Aside from a few pine trees, grasses and gramineous weeds, there is no arable land on BH. Yet, to rule out the possibility that trapped *E. balteatus* individuals originated on BH itself, intensive field surveys were carried out throughout the monitoring period. Trapping data were used to describe *E. balteatus* population dynamics and to infer its migration patterns.

### Backward trajectory analysis

Backward trajectory analysis is widely used for inferring the movement patterns and population sources of migratory organisms (e.g., Stefanescu et al., 2007; Huestis et al., 2019; Sun et al., 2020). In this study, we used trajectory analysis to identify the possible origin of *E. balteatus* migrants that were trapped on BH island during spring-summer (April to July) and autumn (August to October). First, for trajectory simulation, we arbitrarily selected dates during 2003-2018 in which more than 40 *E. balteatus* individuals were caught (i.e., ‘mass migration events’). A total of 76 ‘mass migration events’ were thus identified, including 42 and 34 in the spring and autumn migration period respectively (**Supplementary file 1 Table S1**). For each of those dates, meteorological data at a 1×1° resolution were extracted -through the Global Data Assimilation System (GDAS)- from the National Oceanic and Atmospheric Administration (NOAA) Air Resources Laboratory.

For each date, (night-time) trajectories were calculated every hour from 18:00 to 07:00 of the subsequent day (i.e., operating time of the BH searchlight trap). When running the NOAA Hybrid Single Particle Lagrangian Integrated Trajectory (HYSPLIT) simulation model in MeteoInfo software (version 1.3.3) (Wang, 2014; Stein et al., 2015), the BH light-trap location was set as the end location. Based upon *E. balteatus* radar recordings (Wotton et al., 2019), simulations were performed for five flight heights i.e., 150, 300, 500, 800 and 1 000 m above sea level. The *E. balteatus* flight speed was set identical to the wind speed, while flight duration was set to 24 h. Trajectory analyses thus yielded migration endpoints i.e., take-off locations and potential source areas. Using ArcGIS software, we equally calculated the percentage of trajectories from a given region while omitting endpoints that fell into large water bodies.

### Hydrogen Isotope Analysis

To track animal migration movements, naturally occurring stable isotopes of various elements (e.g., H, C, N) are efficiently used as endogenous markers For migratory insects including hoverflies, the hydrogen isotope deuterium (δD) has been successfully applied (e.g. Wassenaar and Hobson, 1999; Raymond et al., 2014). In our study, we used δD isoscape ratios to pinpoint the geographical origin of migratory *E. balteatus*.

*Hoverfly sampling.* A total of 869 *E. balteatus* adults were collected. These included 286 individuals captured in the BH searchlight trap between April and October 2014-2018, and 583 individuals obtained through sweep-net sampling during April-October 2017-2018 at different sites across China. Sites were located in the Southwestern Region (SW; 1 site), Middle-Lower Yangtze Plain (YzP; 4), Northern Region (NP; 6) and Northeastern Region (NE; 3) (**Supplementary file 1 Table S2**). Each sample was preserved at -20°C and kept at the Institute for Plant Protection, CAAS in Beijing (China) until laboratory processing.

### Stable isotope recordings

As wing tissue is not part of the active metabolism after adult eclosion, it is generally used for isotope analyses (Wassenaar & Hobson, 1998). Accordingly, the wings of all *E. balteatus* specimens were removed with dissection scissors and subsequently sent to the Stable Isotope Mass Spectrometry Facility, Chinese Academy of Forestry (Beijing, China) for hydrogen isotope (δD) measurements as per Zeng et al. (2019). In brief, syrphid wings were cleaned with a methanol-chloroform solution (1:2) and air-dried overnight. Next, the hydrogen isotope ratio (^2^H:^1^H) of the combusted wings was measured using a Flash EA 1112 HT Elemental Analyzer (Thermo Fisher Scientific, Inc., USA) and Isotope Ratio and Mass Spectrometer (Delta V Advantage IRMS, Thermo Fisher Scientific, Inc., USA). Calculations were done using the formula δ^2^H‰= (R_sample_/R_standard_−1) × 1000 in which R is the abundance ratio of heavy isotope to light isotope, namely ^2^H/^1^H. The laboratory error was estimated to be ±2 ‰. Results are expressed in typical delta (δD) notation, in units of per mil (‰), and the relative standard of δ^2^H was the Vienna Standard Mean Ocean Water (VSMOW).

### Population genetics studies

Population genetics studies are routinely used to assess adaptive capacity of different organisms including insects; the resulting data equally reveal migratory movements within an eco-evolutionary perspective (Kim and Sappington, 2013). In this study, we described *E. balteatus* genetic diversity and population structure using one mitochondrial gene and two nuclear genes.

### Specimen Collections and DNA preparation

From April to October 2017-2018, a total of 670 *E. balteatus* adults were collected. These included 133 long-distance migrants that were caught in BH searchlight traps, and 537 individuals collected using sweep-net sampling at 16 sites (**Supplementary file 1 Table S2**). As above, sampling sites for the field-collected individuals were located within five geographical regions: Southwestern Region (SW; 1), Middle-Lower Yangtze Plain (YzP; 4), Northern Region (NP; 6); Northeastern Region (NE; 4) and Northwestern Region (NW; 1), covering three climatic regions i.e., mid-temperate zone (MTZ), warm temperate zone (WTZ), and subtropical zone (SZ). All samples were preserved at -20°C and kept at the Plant Protection Institute (CAAS, Beijing) until further processing. From each specimen, the total genomic DNA was isolated and extracted using a DNeasy Blood and Tissue Kit (Qiagen, Hilden, Germany), following the manufacturer’s instructions. The extracted DNA was re-suspended in 80 µl distilled water, and either used immediately or stored at -20°C for subsequent polymerase chain reaction (PCR) analysis.

### PCR Amplification and Sequencing

For population genetic analyses, one partial mitochondrial (mt) gene [cytochrome b (*CYTB*)] and two nuclear genes [18S rRNA, 28S rRNA] were chosen as molecular markers. Primers were used as previously described (Simmons and Weller, 2001; Mengual et al., 2015), and synthesized by Sangon Biotech Co., Ltd. (Shanghai, China). All fragment amplification was performed in a 25-µl polymerase chain reaction (PCR) volume, using 2×GoldStar Master Mix (CWBIO, Beijing, China). Thermocycling conditions were 10 min at 95°C; followed by 35 cycles of 1 min at 95 °C, 1 min at 55°C (COI) or 56°C (Cytb), and 45 s at 72 °C, and a final extension of 72 °C for 10 min. After verification through 2% gel electrophoresis, the resulting PCR products with the correct target size were sent to Beijing Genomics Institute (BGI) Co., Ltd (Beijing, China) for sequencing in both directions.

### Genetic diversity and population genetic structure

Sequencing results were manually edited, checked and assembled with Chromas 2.31 (Technelysium, Helensvale, Australia) and Seqman within the Lasergene suite version 7.1.2 (DNASTAR, Inc., USA). The corrected nucleotide sequences were then aligned using the ClustalW algorithm implemented in MEGA 6.0 with default parameters (Tamura et al., 2013). To assess *E. balteatus* genetic diversity, the following parameters were calculated for the entire dataset and each individual population using DNASP 6.0 (Rozas et al., 2017): number of polymorphic sites (S), number of haplotypes (h), haplotype diversity (*Hd*), nucleotide diversity (Pi), and average number of nucleotide differences (K). Geographical distribution profiles, phylogenetic trees and haplotype networks were also used to visualize the genetic linkages of different sub-populations. Phylogenetic trees were constructed using the maximum likelihood (ML) method with 1000 bootstrap replicates in MEGA 6.0 (Tamura et al., 2013), while haplotype networks were built in Network 4.6 with the median-joining algorithm (Bandelt et al. 1999). To further evaluate the degree of population differentiation, an analysis of molecular variance (AMOVA) and pairwise population differentiation (F*_ST_*) were carried out using Arlequin 3.059 with 10,000 random permutations (Excoffier and Lischer, 2010). Genetic distances between sub-populations were calculated with a coalescent-based approach using the Bayesian search strategy implemented in MIGRATE-N v. 3.2.1 (Beerli, 2006). To infer demographic history, we used two neutrality tests i.e., Tajima’s D (Tajima, 1989) and Fu’s *Fs* (Fu, 1997), and mismatch distribution analysis (Rogers and Harpending, 1992). Tajima’s D and Fu’s *F_S_* are expected to be nearly zero in an effective population of constant size, with negative or positive values indicative of a respective population expansion or recent bottleneck (Fu, 1997). Populations at demographic equilibrium exhibit a multimodal mismatched distribution, while unimodal patterns reflect recent demographic or area expansions (Rogers and Harpending, 1992). All analyses were conducted using Arlequin 3.5 (Excoffier and Lischer, 2010).

### Pollen grain analysis

Pollen grains carried by insects are routinely used to assess feeding history or movement patterns (e.g., Jones and Jones, 2001). In this study, we used pollen grains that adhered to the body of migrating *E. balteatus* adults and integrated morphologically-based approaches through Scanning Electron Microscopy (SEM) and molecular tactics based upon DNA barcoding. This twin method has been previously employed in our laboratory (e.g., Liu et al, 2016).

### Sample collection, pollen preparation and SEM examination

Over 2014-2018, several sub-samples of 20 migratory *E. balteatus* (or all individuals if the total capture size was below 20) were randomly taken from the BH searchlight trap sample. As such, a total of 1,014 individuals were obtained (**Supplementary file 1 Table S3**). First, all collected samples were examined at 200× magnification using a stereomicroscope (Olympus SZX16, Pittsburgh, PA, USA). Next, suspected pollen grains were gently removed from the insect body and mounted on aluminum stubs with double-sided sticky tape. Next, pollen samples were sputter-coated with gold, and visualized with a Hitachi S-8010 cold field emission scanning electron microscope (Hitachi, Tokyo, Japan) at the Electronic Microscopy Centre of the Institute of Food Science and Technology (CAAS, Beijing, China), or a Zeiss Field Emission scanning electron microscope (Merlin, Zeiss, Germany) at the National Center for Electron Microscopy and School of Materials Science and Engineering, Tsinghua University, Beijing.

### Molecular analysis of single pollen grains

Genomic DNA was extracted from single pollen grains using protocols adapted from Chen et al. (2008). In brief: pollen grains were transferred to individual PCR tubes that contained 5 µL of lysis solution (0.1 M NaOH plus 2% Tween-20), and incubated for 17 min 30 s at 95 °C in a thermocycler (GeneAmp PCR System 9700, Applied Biosystems, Foster City, CA, USA). For each lysis solution, 5 µL Tris-EDTA (TE) buffer was added and the resulting solution was used as a template for subsequent PCR amplifications. To improve species-level identification, four DNA barcoding loci for plants were used simultaneously i.e., two mitochondrial spacer elements ITS1 and ITS2, and chloroplast *rbcL* (Fay et al., 1997; Fazekas et al., 2008; Cheng et al., 2016). All partial regions were separately amplified using DreamTaq DNA polymerase (Thermo Fisher Scientific, Waltham, MA) with the following conditions: an initial denaturation step (95°C for 3 min), followed by 38 cycles at 95°C for 1 min, 55°C for 30 s, 72°C for 1 min, and a final extension of 10 min at 72°C. The resulting PCR products were gel-purified with a Gel Extraction Kit (TransGen, Beijing, China) and ligated directly into the pClone007 Vector (Tsingke, Beijing, China). Positive clones were randomly selected and sequenced with M13 primers using Sanger sequencing at Sangon Biotech Co., Ltd (Shanghai, China) or Tsingke Biotechnology Co., Ltd (Beijing, China).

### Pollen and plant host identification

Each pollen grain was identified based upon its molecular and morphological characteristics, and geographic distribution. First, using the online BLASTn search program, genetic sequences were compared with those at the National Center for Biotechnology Information (NCBI) database. If the sequence top bit score matched with a single species, multiple species within a given genus or multiple genera within a given family, then the sequence was designated to the respective species, genus or family. Sequences that aligned with multiple families were termed to be ‘unidentifiable’ (Hawkins et al., 2015). As such, several sequence taxa were assigned to the rank of genus or family. Separate analyses were performed for the four tested markers and results were combined to identify a given pollen species. Identifications based upon molecular data were further complemented by morphological characterization, using published SEM images of pollen grains of Chinese flora (Ma et al., 1999; Li et al., 2011) or online search engines and palynological databases (https://www.paldat.org/). Finally, species-level identifications were checked against the Flora of China Species Library (https://species.sciencereading.cn) and Plant Science Data Center (https://www.plantplus.cn/cn), to determine the presence of plant hosts within the broader study area.

### Molecular gut content analysis

Molecular gut content analysis has been successfully used to clarify trophic relationships such as herbivory, predation or parasitism (e.g. Staudacher et al., 2011). In this study, laboratory assays were conducted to identify the plant hosts of pollinivorous *E. balteatus*.

Specifically, experimental feeding assays were run to quantitatively assess identification accuracy and detectability half-lives for plant DNA through diagnostic or quantitative real-time PCR (qPCR).

### Insect rearing and pollen collection

For this experiment, *E. balteatus* individuals were sourced from a laboratory population. A laboratory colony was established from individuals sourced in wheat fields at the CAAS Experimental Station in LangFang (LF; Hebei, China; 39.53°N, 116.70°E) during May 2017, and maintained at 25 ± 1 °C, 65 ± 5% relative humidity (RH), and a 16:8 L:D photoperiod. Hatched *E. balteatus* larvae were fed on *Megoura japonica* aphids, and adult flies were fed 10% (w/v) honey solution and pollen (Li et al., 2021). As experimental food in feeding assays, pollen was provided of three preferred host plants: *Cannabis sativa, Humulus scandens* and *Helianthus annuus*. This pollen was directly collected from flowering plants near LF experimental fields during autumn of 2018. After collection, pollen of a given plant species was stored separately in a sterile recipient and kept refrigerated at 4 °C.

### Feeding assay

Newly emerged *E. balteatus* adults were held individually in a disposable plastic cup (30×30×30 cm), provided with a 10% (wt⁄wt) sucrose solution for one day, and subsequently starved for 12 h. Access to water was ensured through a saturated cotton wick. Next, healthy and active hoverfly adults were randomly assigned to either of three diets. More specifically, a given *E. balteatus* adult was individualized in (10 cm diam., 2.6 cm high) plastic Petri dishes containing either of target pollen species i.e., *C. sativa, or H. scandens* or *H. annuus*. Assays were run at room temperature. Over the span of one hour, individuals were observed every 5 min to confirm feeding. Next, those individuals that had fed at least 15 min were selected and transferred to a plastic cup. In this cup, *E. balteatus* adults were given access to a moistened cotton ball (water only) and kept for 0, 2, 4, 8, 16, 24,32, 48 and 72 h. For each treatment, a minimum of 8 samples (replicates) were obtained and individual adults were freeze-killed immediately at each time point. Next, each hoverfly adult was individualized within 1.5-ml tubes and stored at -20 °C for subsequent molecular assay. Some adults were not allowed to feed and were freeze-killed at 0 hr – thus serving as negative controls.

### DNA Extraction

Total DNA was extracted from dissected abdomens with the DNeasy Blood and Tissue Kit (Qiagen, Hilden, Germany), according to the manufacturer’s recommendations. DNA extracts were normalized to 20 ng/µL using Qubit fluorimeter quantification (Invitrogen), and either used directly for PCR amplification or stored at -20 °C until further processing. To avoid plant DNA contamination from the hoverfly’s outer body surface, a bleaching method was adapted from earlier studies and used prior to DNA extraction ( Wallinger et al. 2013). In brief: each individual was washed with 1.5% NaOCl (Beijing Chemical works, Beijing, China) for 10 s and then rinsed with molecular analysis-grade water. Preliminary trials showed that this removed plant DNA contamination from the body surface without destroying the ingested DNA in *E. balteatus* guts. To eliminate any further environmental contamination, all working spaces and equipment were regularly cleaned with 70% ethanol.

### PCR Amplification and Sequencing

To identify pollen loads or host plants from insects, ITS2 offers a high success rate, enables subsequent amplicon sequencing, and counts with a high number of reference sequences among plant DNA barcodes (Han et al., 2013; Pang et al., 2013). Universal ITS2-targeted primers (Cheng et al., 2016) were thus used to specifically detect plant DNA in the feeding assay. PCR assays, TA cloning and sequencing were performed as above. The resultant sequences were identified using BLASTn, and corrected plasmids containing the cloned fragments were used to construct a quantification standard curve.

### Quantitative real-time PCR

For each of the three plant species, we quantified the amount of DNA present in a given *E. balteatus* individual at different times with the respective primer pair through qPCR. Species-specific primers and probes were designed from sequences using the above universal ITS2-targeted primers and synthesized by BGI (Beijing, China) (**Supplementary file 1 Table S4**). qPCRs were performed with the TaqMan method in 20 μl reaction agents composed of 10 μl of 2x QuantiTect Probe PCR Master Mix (Qiagen, Hilden, Germany), 0.5 μM of each primer, 0.2 μM probe, and 1 μl of template DNA, using a 7500 Fast Real-time PCR System (Applied Biosystems). Thermocycling conditions were as follows: 95°C for 2 min, followed by 40 cycles of 94°C for 10 s, 60°C for 10 s. To avoid technical errors, all qPCR reactions were replicated three times. To quantify the amount of DNA of *C. sativa, H. scandens* and *H. annuus*, the respective standard curve equations were used: y = -0.3239x+12.313 (R^2^ = 0.9975), y = -0.3014x+12.372 (R^2^ = 0.9994), and y = -0.3092x+12.505 (R^2^ = 0.9993) (**Supplementary file 1 Figure S4**). In the above equations, y equals to the logarithm of plasmid copy number to base 10, while x = Ct value.

### DNA metabarcoding of gut contents

Metabarcoding has revolutionized species identification and biomonitoring using environmental DNA, offering bright prospects for dietary analyses (Pompanon et al., 2012). In this study, we combined PCR-based gut content analysis with high-throughput sequencing (HTS) of field-collected individuals. The host plant identification protocol could thus be validated under ‘real-world’ conditions and further insights can be gained into *E. balteatus* adult food choice.

### Sampling and DNA extraction

Two sets of adult *E. balteatus* specimens were collected: 180 long-distance migrants captured in the BH searchlight traps from April to October 2015-2018, and 434 individuals collected through sweep-net sampling at 19 sites from April to October 2017-2018. Field-caught populations originated from five geographical regions of China (**Supplementary file 1 Table S2**). DNA from field-caught samples was individually extracted using the DNeasy Blood and Tissue Kit (Qiagen, Hilden, Germany), and subject to experimental procedures as described above.

### MiSeq sequencing of ITS2 barcode gene amplicons

A two-step laboratory protocol was followed: the ITS2 fragment was first amplified using the barcoded universal primers to generate mixed amplicons. Next, amplicons were sequenced using HTS for comparison against known barcode references. In the first step, we used PCR to uniquely index each sample using the modified universal primers that were tagged with a sample-specific eight-mer oligonucleotide tag at the 5′-end. In total, 90 sets of index primers were used to amplify the ITS2 region. This was done to ensure that multiple samples could be processed simultaneously into a single sequencing run and be subsequently separated via bioinformatics processing. Each sample was processed in three independent PCRs to avoid reaction-specific biases. Each of the replicate PCRs consisted of 2 µl of the DNA template, 12.5 µl of 2×GoldStar Master Mix (CWBIO, Beijing, China), and 0.5 µM of amplicon primers in a 25 µl reaction volume with the following PCR programme: 95°C for 10 min, 38 cycles of denaturation at 95 °C for 1 min , annealing at 49°C for 40 s and elongation at 72°C for 40 s, and a final extension step at 72°C for 10 min. After reaction, the replicate PCRs for each sample were combined and gel-purified using the MinElute Gel Extraction Kit (QIAGEN, Hilden, Germany) as per manufacturer instructions and quantified with a NanoDrop ND-2000 UV-vis spectrophotometer (NanoDrop Technologies, Wilmington, DE). Next, purified amplicons with different tags were pooled in an equal concentration to make a composite DNA sample and preceded for HTS sequencing. A negative control with no DNA template was run in parallel. All pooled amplicons (up to 90 specimens each) were sent to the GENEWIZ, Inc. (Suzhou, China) for final amplicon library construction and Illumina HTS. Briefly, for each mixed-amplicon, the indexed Illumina-compatible libraries were constructed using VAHTS Universal DNA Library Prep Kit for IlluminaV3 (Vazyme Biotech Co., Nanjing, China) with standard protocols, and subsequently sequenced on the Illumina MiSeq platform (Illumina, San Diego, CA, USA) using a 2x300bp paired-end (PE) configuration, as per manufacturer protocols.

### Data analysis

Paired-end reads were assigned to samples according to the unique 8bp barcode of each sample and truncated by cutting off the barcode and primer sequence, reads containing ploy-N, and low-quality reads. The trimmed forward and reverse reads were joined based on overlapping regions within paired-end reads. Next, chimeric sequences were removed after being identified by comparing merged sequences with the reference RDP Gold database using UCHIME algorithm. The remaining high-quality reads were then clustered into Operational Taxonomic Units (OTUs) at a 99% sequence similarity level, using VSEARCH1.9.6. Representative OTU sequences were taxonomically annotated with BLASTn searches against the NCBI database (accessed 02/2021) using the remote command line interface (Camacho et al., 2009). For each sequence, the top 50 matches were written to a tsv file, but matches were only included in downstream analyses once they met the following criteria: 1) min. 90% of the query sequence is present in the BLASTn-generated subject sequence (sequence coverage), 2) at least 95% sequence similarity (identity), 3) an e-value below 0.001, and 4) a subject sequence derived from ITS2 of a land plant (Prosser and Hebert, 2017). All samples that yielded a sequence that met these thresholds from one or both PCR replicates were considered as host identifications. Annotation results were analyzed following Hawkins et al. (2015). Furthermore, all BLAST results were verified using expert knowledge, integrating an understanding of local habitats, species distribution and rarity to refine plant host identifications. For each dietary item, occurrence frequency was calculated as the number of samples in which the item was present divided by the total number of samples. Also, the relative read abundance was calculated by dividing the number of reads of each dietary item (and individual) by the total number of reads in the sample.

### Statistical Analysis

Differences in *E. balteatus* trap capture rate were analyzed using a zero-inflated generalized linear model (GLM) (Brooks et al., 2017). To compare counts among years, post-hoc tests were run using the emmeans package (Searle et al., 1980). Wilcoxon rank-sum test was used to compare differences in the δD values of migratory *E. balteatus* wings sampled at different times. A chi-square test was used to compare the differences in the frequency of pollen deposits on *E. balteatus* during different migration phases and the characteristics of pollen source plants. To compare δD and alpha diversity values between groups, we used One-way analysis of variance (ANOVA) followed by Tukey’s test for multiple comparisons. Venn and upset diagrams were drawn to represent the interactions among host plant communities of different groups using the omicshare cloud tool under default instructions (http://www.omicshare.com/). Principal coordinate analysis (PCoA) and Analysis of similarities (ANOSIM) was used to evaluate differences in *E. balteatus* host plant abundance between specific migration phases or regions. To identify features characteristic of certain groups, we built a random forest machine learning model based on genus abundance data from 180 *E. balteatus* migrants captured on BH. All statistical analyses were performed in R version 4.0.3 (R Development Core Team, 2020).

## Results

### Migration dynamics

Despite the local availability of weedy host plants, intensive field surveys did not detect a presence of *E. balteatus* larvae on Beihuang (BH) island. Yet, night-time trapping consistently yielded *E. balteatus* adults from late April to October (**Figure 1A**) throughout the 16-yr sampling period. Hoverfly abundance (or trap capture rate) showed important annual variation (Marginal R^2^ = 0.163, χ^2^ = 75.4, *df* = 15, *P* < 0.001) (**Supplementary file 1 Figure S1**), with peak population sizes of 5,068 and 2,381 individuals in 2009 and 2010 respectively. Overall, *E. balteatus* trap capture rate exhibited a bimodal pattern with peak abundance from May-June and August-September, thus comprising two distinct migration stages. On an annual basis, *E. balteatus* migration covered a period of 151.2 ± 17.9 days (Supplementary file 1 **Table S5**).

**Figure 1.**
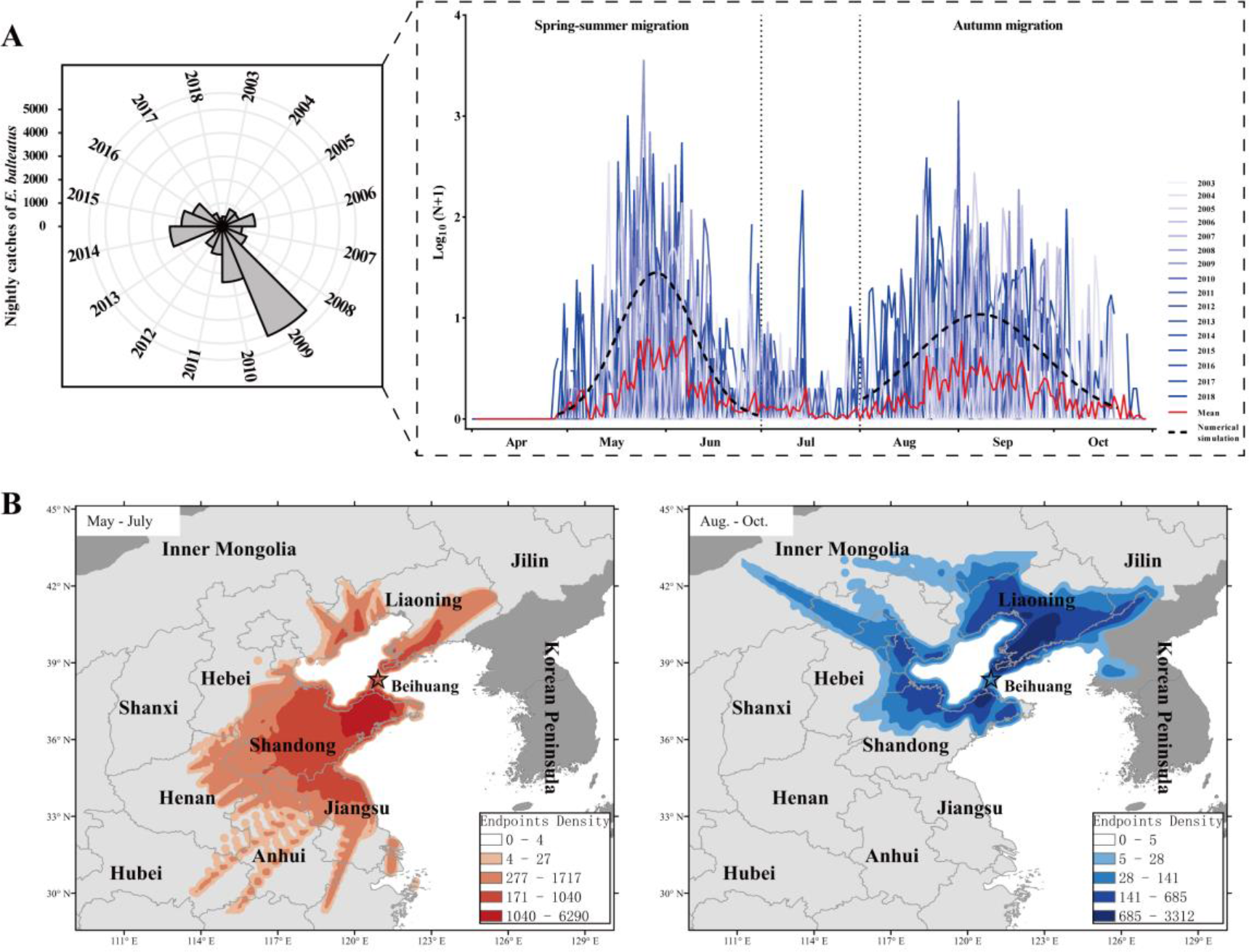
**Seasonal migration patterns of *Episyrphus balteatus* in Eastern China**. **A**, Annual migration dynamics, expressed as nightly searchlight-trap catches, of *E. balteatus* on Beihuang Island (BH; Bohai Gulf, China) from April to October 2003–2018. **B**, Endpoints of backward trajectories of trapped E. balteatus individuals during mass migration events over 2003–2018 for a 24 h flight duration. Darker colors indicate a higher density of endpoints at a particular location. The left panel (i.e., orange colors) reveals the possible source areas of late spring and summer immigrants, while the right panel (i.e., blue colors) indicates those of autumn immigrants.

Next, the possible origin of *E. balteatus* migrants on BH was identified through backward trajectory analysis using the HYSPLIT model. For ‘mass migration’ events over 2003-2018 (**Supplementary file 1 Table S1**), spring-summer migrants primarily originated in southern areas, while autumn migrants arrived from nearby areas in any direction (**Figure 1B**). During spring-summer, more than 92% endpoints where located south of BH i.e., in Shandong, Jiangsu, Henan and Anhui provinces; during autumn, 73% endpoints were distributed in northern areas.

To corroborate the above patterns, wings of BH migrants and wild-caught individuals from sites across China were subject to hydrogen isotope analysis. On BH, greater variability in δD values was recorded for autumn versus spring-summer migrants over the entire study period (Wilcoxon rank sum test; W = 11.93, df = 254.13, p <0.0001) (**Figure 2B**). By comparing the above δD values with its established precipitation gradient, source areas of *E. balteatus* migrants were identified. Considering how δD values in field-collected populations divert from latitudinal gradients (**Figure 2C**), seasonal differences became apparent upon a geographical grouping of *E. balteatus* populations. Overall, spring-caught adults showed higher δD values than those captured during autumn (Wilcoxon rank sum test; NE subgroup: W = 3232.5, p< 0.001; NP: W = 13804, p< 0.001; YzP: W = 3440, p< 0.001) (**Figure 2D**).

**Figure 2.**
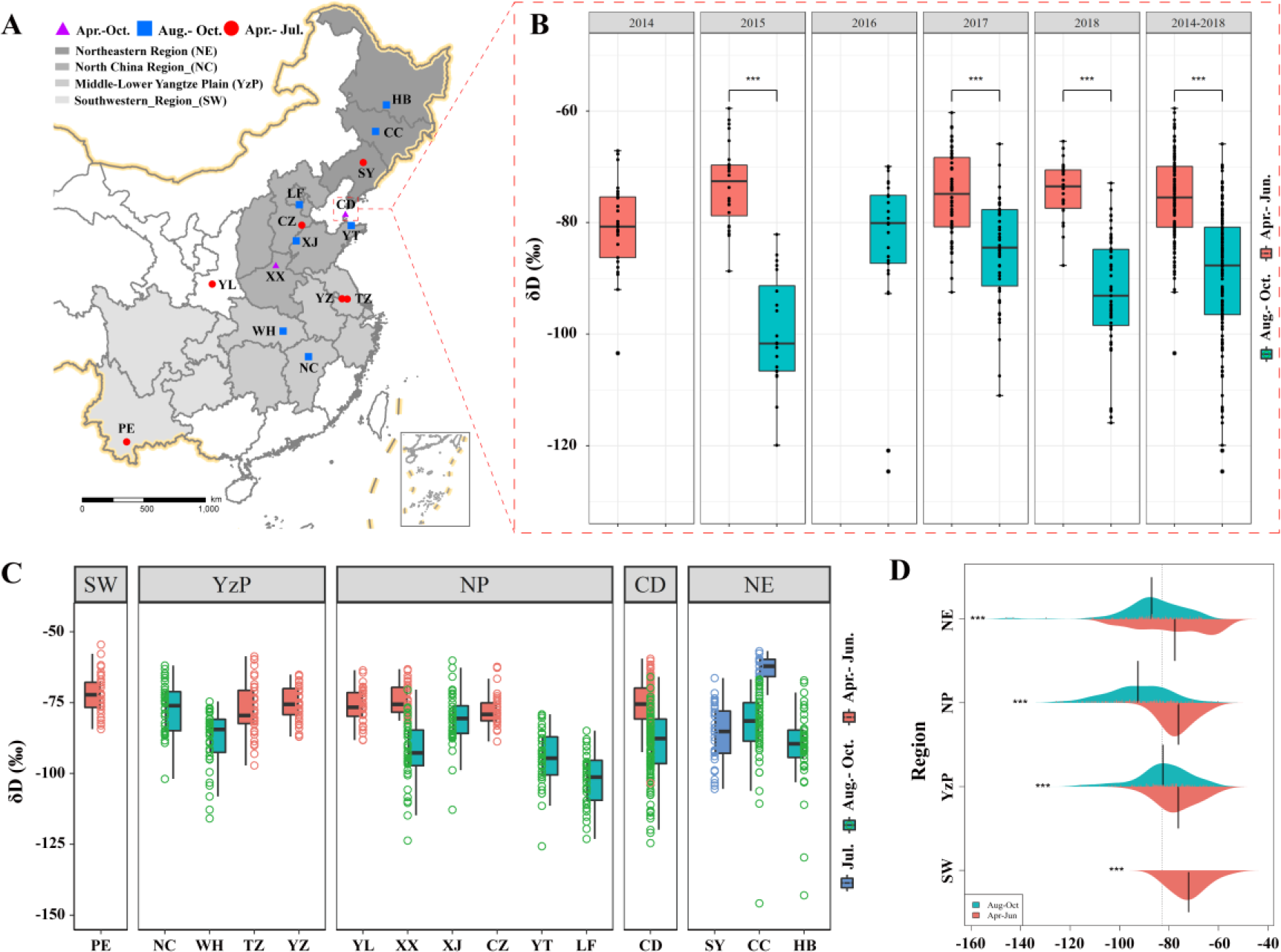
**Stable hydrogen isotope analysis of *E. balteatus***. **A**, Map of the the sampling area. **B**, Seasonal incidence of mean δD values in the wings of migratory *E. balteatus* adults, as recorded for different migration stages over 2014-2018 on Beihuan Island (China). Double asterisks (**) indicate a statistically significant difference (p < 0.01, t test). **C-D**, Seasonal incidence of mean δD values in the wings of wild-caught individuals collected in different parts of China from April to October 2017-2018. Locations are presented in order of their geographical position along a longitudinal gradient.

Isotope patterns reflected how hoverfly adults engage in bi-directional migration over hundreds of kilometers.

### Population genetics

To gain genetic evidence of its regional migration, we described *E. balteatus* genetic diversity and population structure using one mitochondrial DNA gene (Cytb) and two nuclear DNA genes (i.e., 18s rRNA, 28s rRNA). Upon analysis of 670 field-collected specimens and 133 light-trapped individuals (representing a respective 16 and 2 populations), high haplotype diversity and low nucleotide diversity was recorded (**Table 1**). Based upon Cytb sequences, 83 haplotypes were identified among 530 individuals, with haplotype diversity (*Hd*) ranging from 0.0800 (XJ) to 0.857 (SH) (total= 0.351) and nucleotide diversity (π) from 0.000110 (XJ) to 0.00536 (LF) (total= 0.00178), respectively. Conversely, the concatenated nuclear gene possesses improbably high haplotype diversity and low nucleotide diversity. Up to 145 haplotypes were detected among 260 individuals with *Hd* ranging from 0.810 (HB) to 1 (YP, YL and PE) (total=0.961) and π ranging from 0.00392 (XX) to 0.01384 (CDI) (total=0.000008).

**Table 1.** Genetic diversity indices of 18 E. balteatus populations based on Cytb and 18S-28S rRNA gene. . n, sample size; S, number of segregating sites; H, number of haplotypes; Hd, haplotype diversity; K, average number of differences; Pi, nucleotide diversity.

|  | CytB |  |  |  |  |  |  | 18S-28S rRNA |  |  |  |  |  |  |
| --- | --- | --- | --- | --- | --- | --- | --- | --- | --- | --- | --- | --- | --- | --- |
|  | n | S | H | Hd | K | Pi | PiJC | n | S | H | Hd | K | Pi | PiJC |
| LF | 23 | 37 | 7 | 0.46245 | 3.92095 | 0.00536 | 0.00543 | 12 | 25 | 8 | 0.84848 | 5.51515 | 0.00452 | 0.00454 |
| XX | 42 | 6 | 5 | 0.33682 | 0.45296 | 0.00062 | 0.00062 | 16 | 31 | 10 | 0.825 | 4.79167 | 0.00392 | 0.00394 |
| HB | 23 | 1 | 2 | 0.08696 | 0.08696 | 0.00012 | 0.00012 | 18 | 28 | 10 | 0.81046 | 5.60784 | 0.00459 | 0.00461 |
| WH | 34 | 10 | 5 | 0.2246 | 0.58824 | 0.0008 | 0.00081 | 12 | 32 | 9 | 0.90909 | 6.90909 | 0.00566 | 0.00569 |
| CC | 16 | 28 | 7 | 0.625 | 3.5 | 0.00478 | 0.00482 | 11 | 22 | 10 | 0.98182 | 5.63636 | 0.00462 | 0.00464 |
| NC | 21 | 12 | 5 | 0.35238 | 1.65714 | 0.00226 | 0.00228 | 15 | 45 | 12 | 0.94286 | 9.61905 | 0.00788 | 0.00793 |
| SY | 45 | 16 | 6 | 0.25051 | 0.75354 | 0.00103 | 0.00104 | 21 | 49 | 18 | 0.98095 | 9.2619 | 0.00759 | 0.00764 |
| CF | 27 | 21 | 8 | 0.45869 | 2.02849 | 0.00277 | 0.00279 | 16 | 39 | 16 | 1 | 9.00833 | 0.00738 | 0.00742 |
| XJ | 25 | 1 | 2 | 0.08 | 0.08 | 0.00011 | 0.00011 | 18 | 31 | 13 | 0.95425 | 6.71895 | 0.0055 | 0.00553 |
| YT | 29 | 22 | 5 | 0.31773 | 1.95074 | 0.00266 | 0.00268 | 16 | 34 | 12 | 0.95833 | 7.8 | 0.00639 | 0.00643 |
| YP | 18 | 8 | 3 | 0.21569 | 0.88889 | 0.00121 | 0.00122 | 8 | 38 | 8 | 1 | 12.89286 | 0.01056 | 0.01064 |
| YL | 30 | 4 | 6 | 0.31034 | 0.33103 | 0.00045 | 0.00045 | 14 | 35 | 14 | 1 | 8.56044 | 0.00701 | 0.00705 |
| TZ | 28 | 2 | 3 | 0.14021 | 0.14286 | 0.0002 | 0.0002 | / | / | / | / | / | / | / |
| PE | 18 | 7 | 4 | 0.47059 | 1.79739 | 0.00246 | 0.00247 | 11 | 98 | 11 | 1 | 22.96364 | 0.01881 | 0.01922 |
| YZ | 31 | 4 | 5 | 0.24516 | 0.25806 | 0.00035 | 0.00035 | 10 | 72 | 9 | 0.97778 | 15.62222 | 0.01279 | 0.01309 |
| SH | 7 | 8 | 5 | 0.85714 | 2.28571 | 0.00312 | 0.00313 | / | / | / | / | / | / | / |
| CDI | 52 | 43 | 16 | 0.52413 | 2.30166 | 0.00314 | 0.00316 | 25 | 103 | 21 | 0.97667 | 16.9 | 0.01384 | 0.01413 |
| CDII | 61 | 48 | 17 | 0.48251 | 1.70055 | 0.00232 | 0.00234 | 37 | 65 | 28 | 0.98198 | 9.15165 | 0.0075 | 0.00754 |

Though phylogenetic analyses showed four and five distinct clades among the Cytb and nuclear haplotypes, haplotype and geographical origins were not linked (**Figure 3A**). Median-joining also did not reveal geographical clustering. Instead, a star-like pattern was displayed with the most common, ancient haplotypes in the center (**Supplementary file 1 Figure S2**). Most haplotypes were unique to individuals and populations, while only 9 (out of 83 Cytb haplotypes) and 30 (out of 145 nuclear haplotypes) were shared. In each population, shared haplotypes occurred at 63-100% frequencies for *Cytb* and 33-88% for nuclear genes (**Figure 3B**). Only 1.8% and 11.1% of genetic variation for the respective *Cytb* and nuclear gene could be attributed to variation among populations (**Supplementary file 1 Table S6**). Low pairwise *F_ST_* values between different localities equally reflected low levels of genetic differentiation (**Figure 3C**). When describing effective population sizes and migration rates between (geographically grouped) populations, high levels of inter-population gene flow were recorded (**Figure 3D**). Migration between *E. balteatus* populations was asymmetrical and exhibited a general migration trend towards the Yangtze basin, with north>centre, south>centre and west>east as dominant directions. Migration thus enables genetic mixing among *E. balteatus* populations from remote origins. Given the negative Tajima’s *D* and Fu’s *Fs* (*Cytb*: Tajima’s *D* = -2.72034, Fu’s *F_S_* = -27.4859; concatenated 18S-28S rRNA: Tajima’s *D* = -2.33710, Fu’s *F_S_* = -34.456) and unimodal distribution for both markers (**Supplementary file 1 Figure S3**), *E. balteatus* populations possibly experienced recent population expansion.

**Figure 3.**
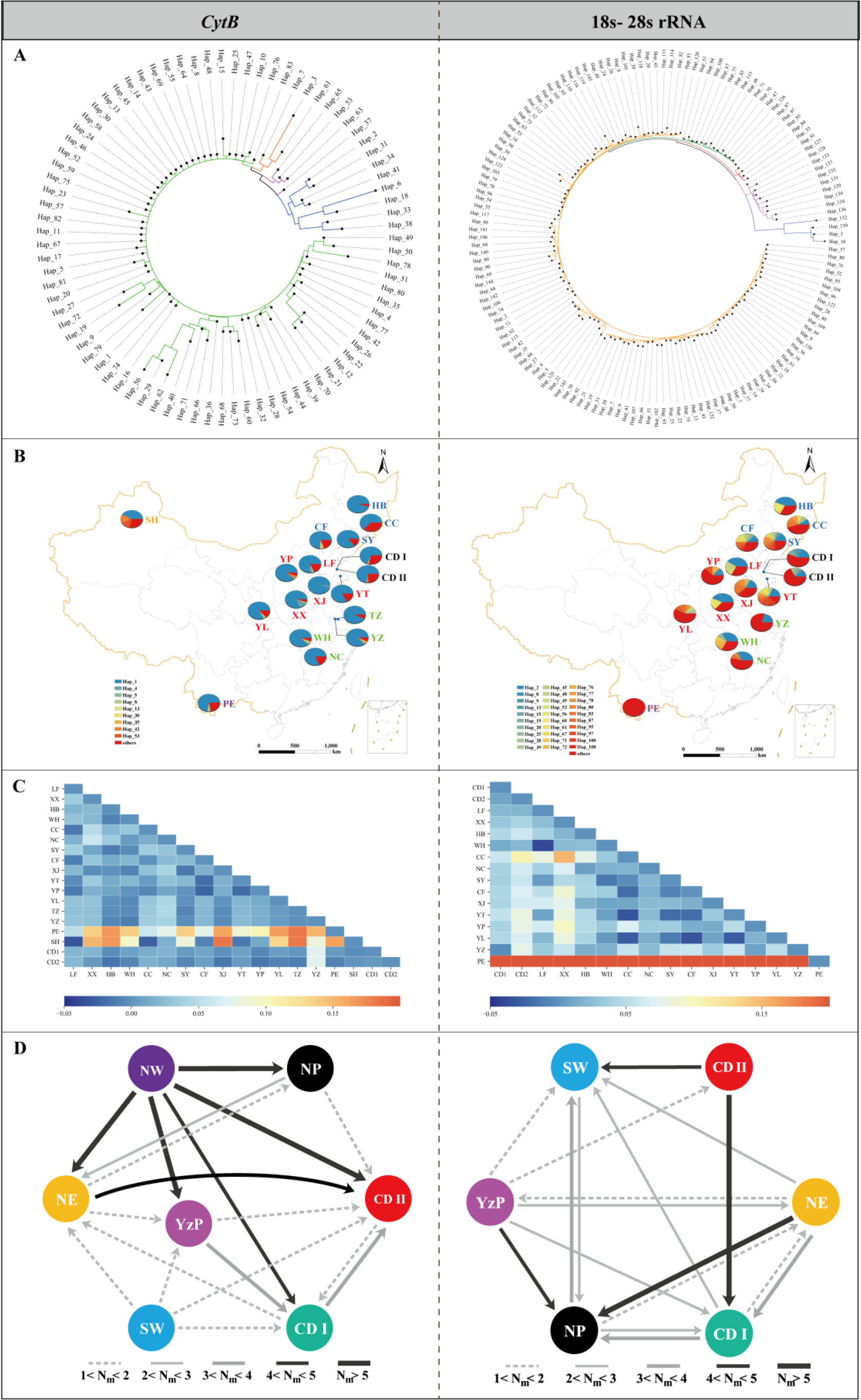
**Population genetic structure of 18 Chinese *E. balteatus* populations**. Analyses are either based on one mitochondrial Cytb gene (left panel) or on combined 18s and 28s rRNA nuclear genes (right panel). A, Neighbor-joining (NJ) phylogenetic trees of haplotypes, in which different clades are depicted with different colors. B, Spatial distribution of E. balteatus haplotypes. At each given location, a pie-chart shows the proportional abundance of haplotypes. C, Heatmap diagram of genetic differentiation coefficient (Fst) between populations. D, Estimated migration dynamics of different populations using MIGRATE-N. Individual arrows represent the prevailing migration direction and the arrow thickness is proportional to the number of migrants. SW, Southwestern Region; YzP, Yangtze Plain; NP, Northern Region and NE, Northeastern Region, as defined in Figure 2.

### Palynological analysis

Morphological characteristics and barcode markers were used to identify pollen species attached to migrating *E. balteatus* adults, and to infer the associated movement and flower visitation patterns. Among 1,014 BH adult migrants collected during 2015-2018, 32% had pollen grains adhering to the body surface. Using DNA sequences, pollen morphology and plant distribution records, 46 pollen species representing at least 42 plant genera and 26 families were identified (**Figure 4, Supplementary file 1 Table S3**). Out of these, 10 were identified to genus level and the remained to species level. Few adults carried pollen from multiple plant species. Pollen-bearing plants mainly pertained to Asteraceae (12), Moraceae (4) and Celastraceae (3), and were primarily herbaceous as compared to woody plants (χ^2^ = 112.26, df = 1, P < 0.001). Highest and lowest pollen adherence ratio levels were recorded in October (50%) and April (17%), respectively, while pollen grain identity equally exhibited seasonal variation (**Supplementary file 1 Table S3**).

**Figure 4.**
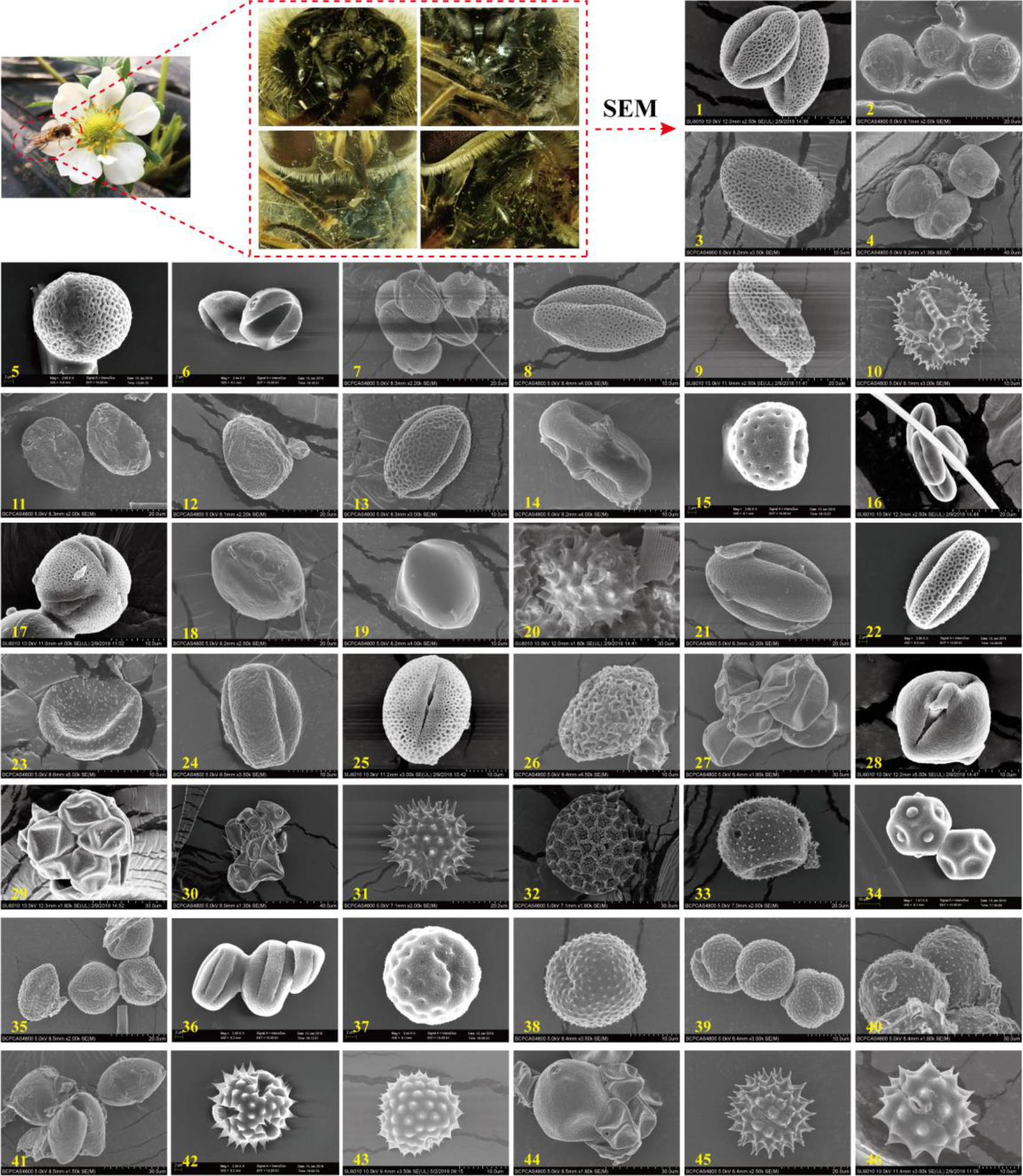
Scanning Electron Microscopy (SEM) microphotographs of pollen grains attached to *E. balteatus* migrants on BH during 2014-2018. 1..*Ailanthus altissima*; 2. *Cotinus coggygria*; 3. *Forsythia suspensa* ; 4. *Prunus avium*; 5. *Brassica* L.; 6.*Morus alba*; 7. *Citrus sinensis*; 8. *Descurainia sophia*; 9. *Euonymus* L.; 10. *Taraxacum* L.; 11. *Sedum japonicum*; 12. *Populus cathayana* ; 13. *Celastrus orbiculatus*; 14. *Daucus carota*; 15. *Chenopodium* L.; 16. *Castanea mollissima*; 17. *Amorpha fruticosa*; 18. *Diospyros lotus*; 19. *Ziziphus jujuba* ; 20. *Cirsium setosum*; 21. *Neoshirakia japonica*; 22. *Flueggea* L.; 23. *Maclura pomifera* ; 24. *Rumex* L. ; 25. *Euonymus* L.; 26. *Schisandra chinensis*; 27. *Eleusine indica*; 28. *Actinidia kolomikta*; 29. *Cannabis sativa* ; 30. *Humulus scandens*; 31. *Helianthus annuus*; 32. *Persicaria orientalis*; 33. *Adenophora trachelioides*; 34. *Gypsophila paniculata*; 35. *Artemisia* L.; 36.*Rubia cordifolia*; 37. *Rubia cordifolia* ; 38. *Artemisia* L.; 39. *Artemisia* L.; 40. *Artemisia* L.; 41. *Allium tuberosum*; 42. *Tripolium vulgare*; 43. *Ambrosia trifida*; 44. *Sorghum bicolor*; 45. *Aster tataricus*; 46. *Chrysanthemum zawadskii*. The scale bar has been showed on the bottom of each photograph.

Specific plant taxa were associated with certain migration stages. Plants endemic to certain ecological zones helped to pinpoint migration origins (Jones and Jones, 2001). During spring-summer, presence of *Citrus* L., *Sedum japonicum* or *Euonymus myrianthus* hinted at migration origins in south-central China. Conversely, during autumn, the presence of *Chrysanthemum zawadskii* and *Adenophora trachelioides* mirrored potential migration origins in northeastern China. These results further reflected northward *E. balteatus* migration flows during spring with return movements in autumn.

### Spatiotemporal dietary shifts

Exploratory assays showed how plant DNA could be reliably detected in hoverfly guts for up to 9d post-feeding (**Supplementary file 1 Figure S5**). By pairing gut content analysis with HTS, (floral) diet profiles of 180 light-trapped *E. balteatus* migrants and 436 field-collected individuals were elucidated. MiSeq paired-end sequencing of the *ITS2* gene yielded 3,348,773 raw reads from 616 samples. After quality filtering and reference-based chimera removal, a total of 2,952,448 sequences remained. These effective reads were clustered into OTUs for further analysis (**Supplementary file 2**). According to the taxonomic assignment, numerically abundant plant species were identified in all populations. Significantly more plant taxa were recorded in the BH migratory population (36 orders, 76 families, 320 genera) as compared to field-collected populations (**Figure 5B**). Overall, 1012 plant species belonging to 39 orders, 91 families and 429 genera were identified, with *Asteraceae* (84), *Poaceae* (37), *Apiaceae* (21) and *Fabaceae* (21) (count at genus level) the dominant families accounting for 38.0% of all plant taxa (**Supplementary file 3**). Further analysis of plant characteristics revealed important similarities with the above palynological analysis, in which more Angiosperm and Dicotyledon plants were recorded than Gymnosperms (χ^2^ = 393.76, df = 1, P < 0.0001) or Monocotyledons χ^2^ = 253.04, df = 1, P < 0.0001). Woody plant hosts were more common than herbaceous ones (χ^2^ = 61.94, df = 1, P < 0.0001) (**Figure 5C**).

**Figure 5.**
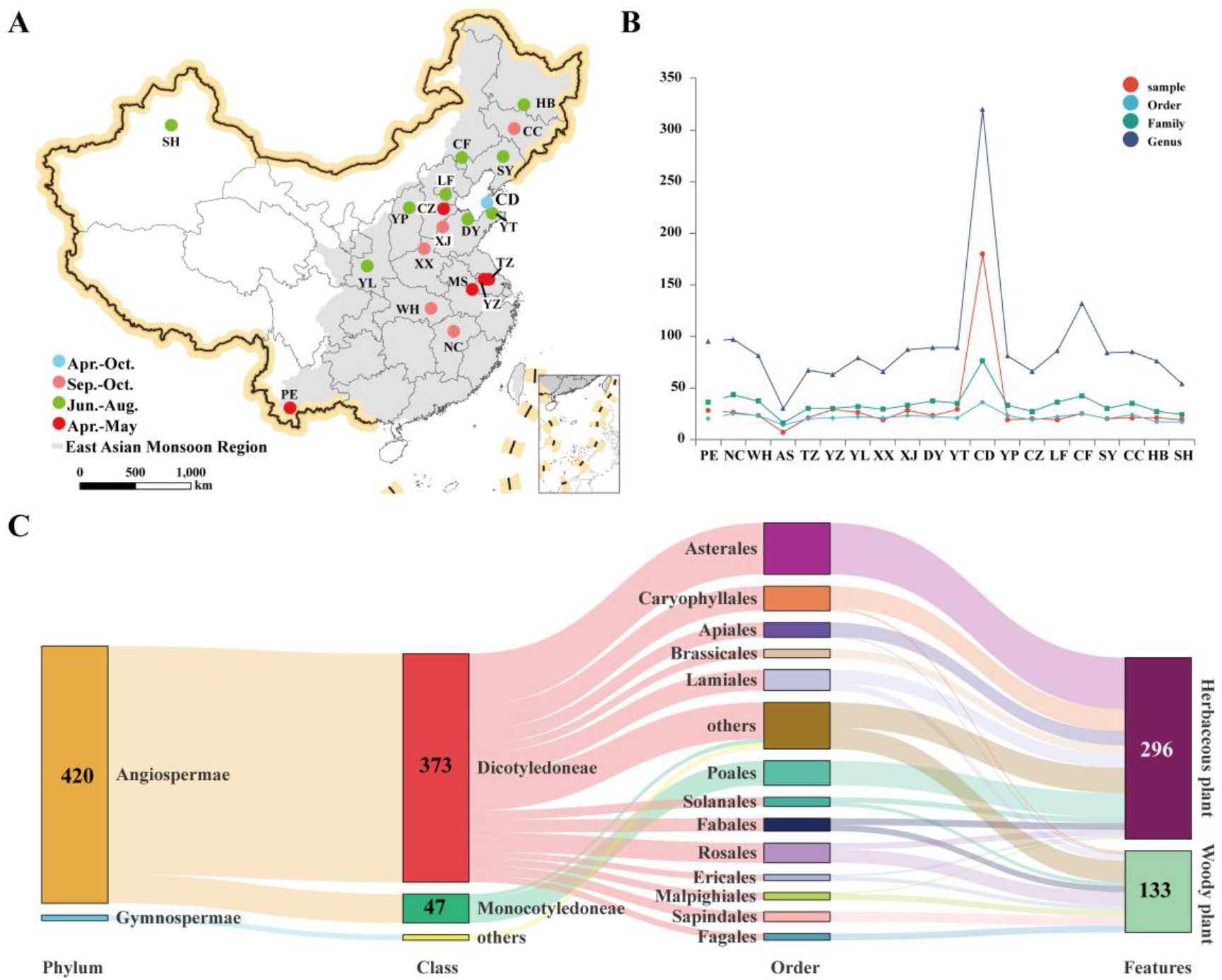
Host plants of *E. balteatus*, as identified through DNA metabarcoding (i.e., for BH migrant individuals) or DNA-based gut content analysis (i.e., for field-collected individuals from 19 sites). A, Geographical location of the sampling sites. B, Taxonomic resolution (i.e., order, family, genus) of the identified host plants for each hoverfly population. C, Taxonomic structure of identified 429 plant genera, with individual numbers referring to the number of counts of plant genera. The detailed information see Supplementary file 3.

While the dominant host plant taxa proved unique for several geographic locations (**Figure 6, Supplementary file 4**) and the relative abundance of the top 10 genera for particular (geographical) populations was very high (**Figure 7A**), plant community structure exhibited weak temporal differences upon a spatiotemporal grouping of *E. balteatus* populations. Venn diagrams and hierarchical cluster analysis indicated that most plant taxa were widespread among locations and sampling times (**Figure 7B-D**). Geographical location thus only affected pollen transport networks to small extent. Importantly, the host plant composition of the BH migratory population almost equals the sum of field-collected populations (**Figure 7D**). Furthermore, plant community composition exhibited important within-year variability. During April-May, *E. balteatus* primarily foraged on *Brassica* spp. (*Brassicaceae*) and *Pinus* spp. Over time, *Gossypium* spp. *and Flueggea* spp. (*Moraceae*), *Ajania* spp. and *Artemisia* spp. (*Asteraceae*) became more common pollen hosts (**Figure 8A,B**). Venn diagram displayed a certain number of unique plant species in all *E. balteatus* groups (**Figure 8C**), with a clear shift in plant taxa over time (**Figure 8D**). Random forest machine learning further revealed the main plant taxa during specific seasons (**Figure 8E**).

**Figure 6.**
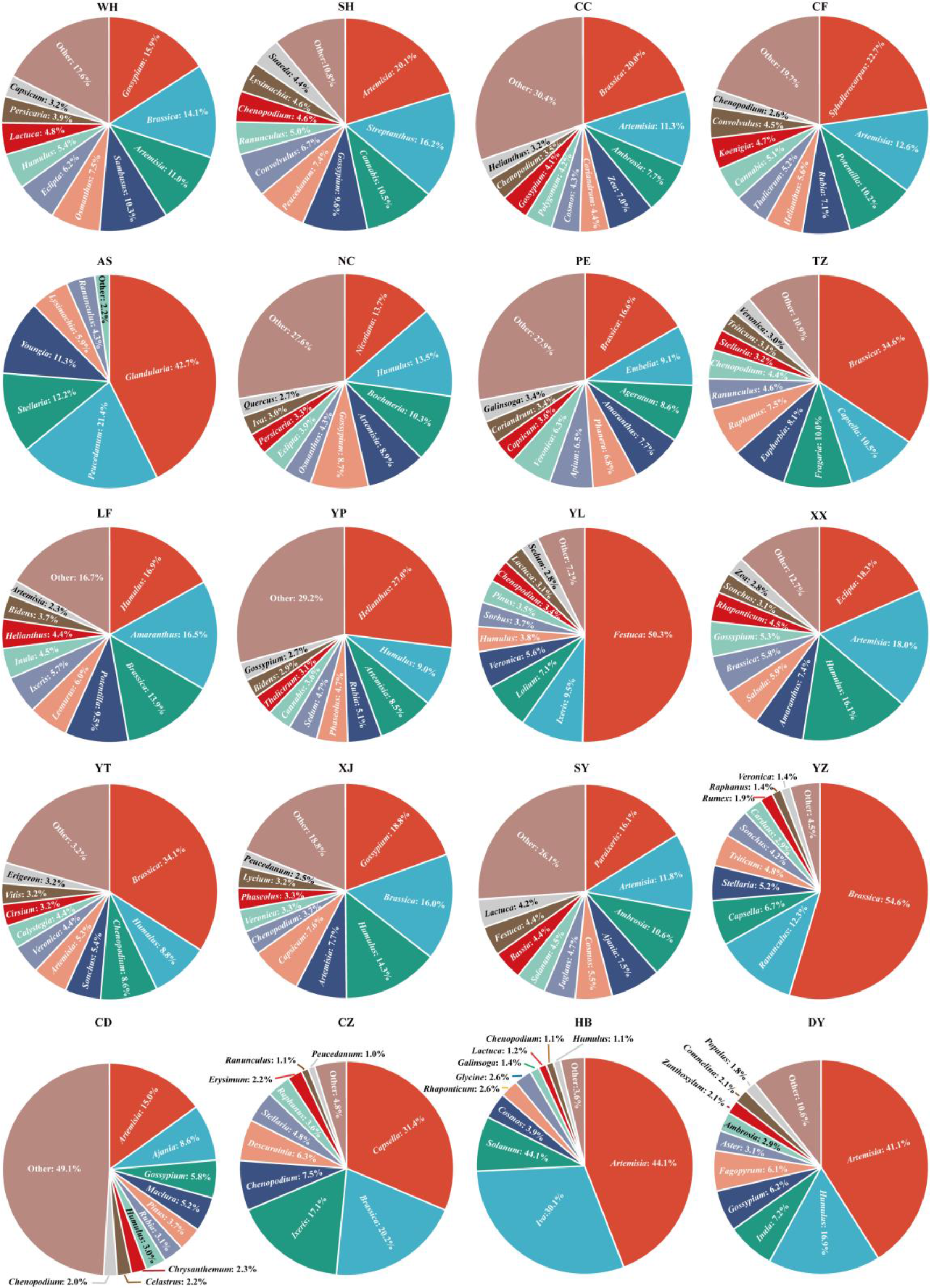
Relative abundance profiles of the 10 prevailing host plants of adult *E. balteatus*. Abundance profiles are drawn per region and migration season. Host plants are identified at the genus level.

**Figure 7.**
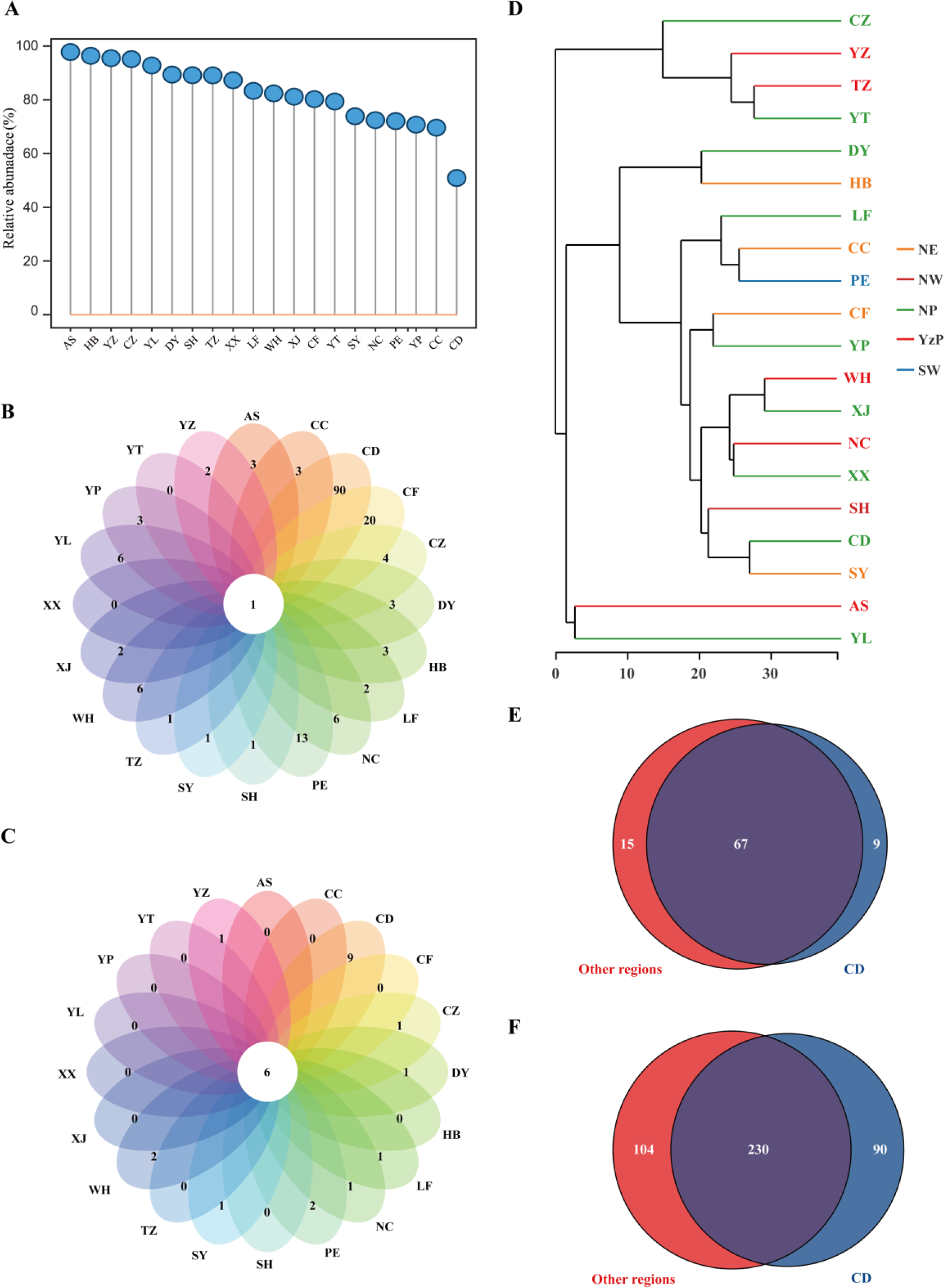
Spatial variability in *E. balteatus* host plant usage at 20 sites across China, as inferred through molecular gut content analysis. A, Spatially-delineated taxonomic distribution of the 10 prevailing host plants. B-C, Venn analysis for host plants across different regions. Plant hosts are specified at the family/genus level. D, Hierarchical clustering tree for plant hosts, as specified at the genus level. E-F, Venn diagram showing the common and unique host plants associated with E. balteatus migrants captured on BH and field-collected individuals. Plant hosts are specified at the family/genus level.

**Figure 8.**
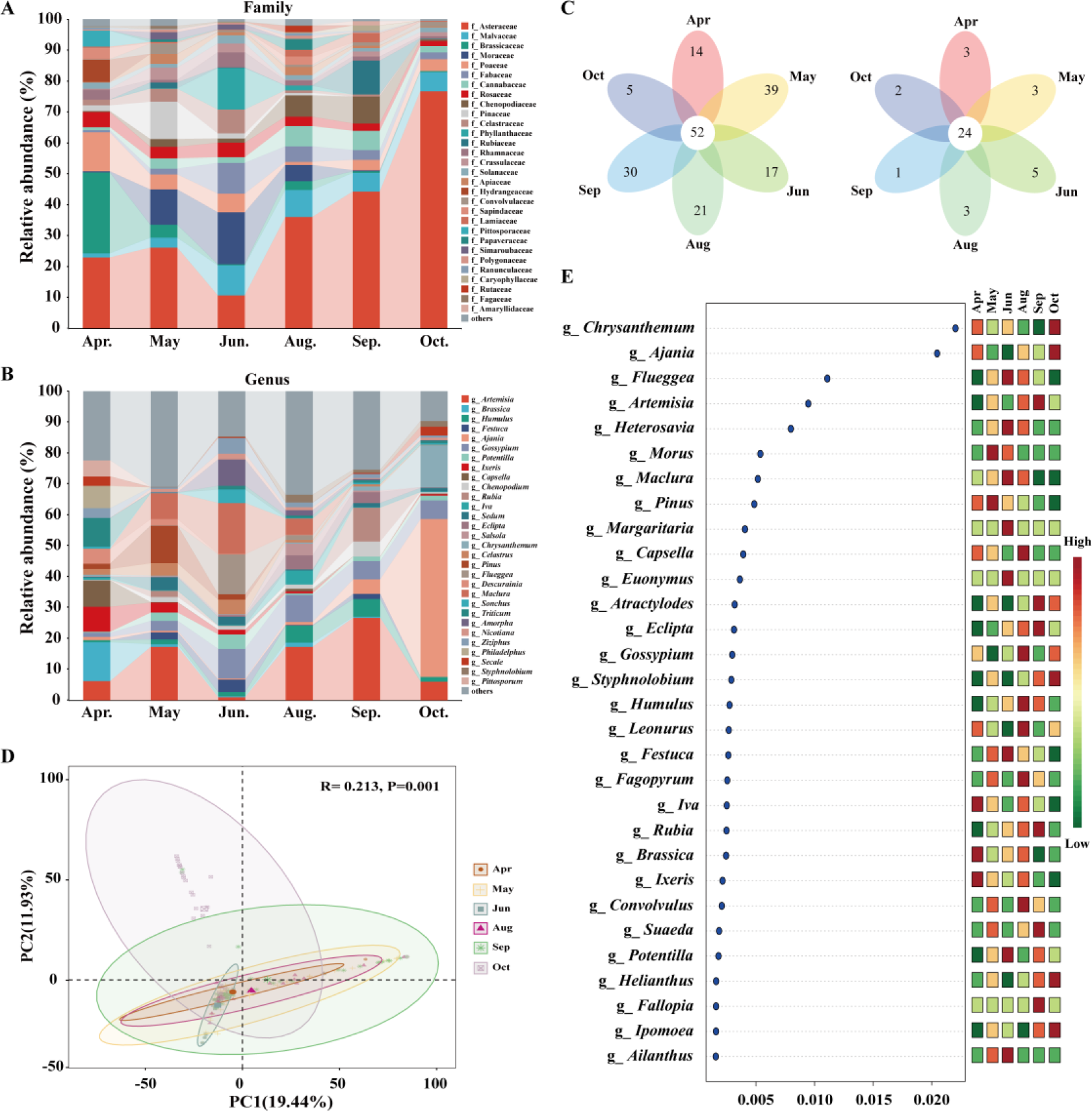
Temporal shifts in host plant usage of *E. balteatus* migrants on BH during different migrating seasons, as determined by molecular gut content analysis. A-B, Relative abundance of the 30 most common plant taxa over time. Plant hosts are listed at the family or genus level. C, Venn analysis for plant hosts during different migration seasons, with plants specified at family or genus level. D, Principal coordinate analysis (PCoA) of the host plants based on the unweighted UniFrac distance metrics on genus level. ANOSIM, R = 0.213 , P = 0.001. E, Top 30 plant genera prioritized by random forest analysis and ranked by the mean decrease in accuracy.

## Discussion

In eastern Asia, BeiHuang (BH) island has lent itself to decipher the migration behavior of more than 100 insect species (Guo et al., 2020). For a widely distributed hoverfly species (*Episyrphus balteatus*) in China, our work unravels the extent to which seasonal migration events mediate ecosystem service delivery. Specifically, multi-year monitoring at BH showed how *E. balteatus* populations undertake cross-maritime migration, moving southward during spring-summer and returning during autumn. Population genetics reveals how *E. balteatus* annually engages in bi-directional migration over hundreds of kilometers and how the species exploits a spatio-temporally variable community of pollen-shedding plants. In view of the unrelenting global environmental degradation and the precipitous decline of insect populations, our work helps guide interventions to conserve this beneficial insect species and to safeguard its vital ecosystem services.

While other hoverfly species in Europe and elsewhere have been shown to undertake long-distance migration (e.g.,Wotton et al., 2019; Gao et al., 2020), this study provides (to our best knowledge) the first evidence for such phenomenon in East Asia. Our findings further refute the prevailing thought that hoverfly species exclusively engage in diurnal migration. As the diurnal *E. balteatus* thus adopts night-time migration in a similar way as certain dragonfly or beetle species (Feng et al., 2006; Anderson, 2009), our work shines new light on the migration strategies of these globally important beneficial insects. As such, certain hoverfly species likely possess specific behavioral traits to undertake both nocturnal and diurnal migration (Michalik et al., 2020). Given the major ecological implications of such ‘dual’ migration strategy e.g., in terms of pollination or pest control, this knowledge gap urgently needs to be filled.

As equally noted for other migratory insects (e.g., Guo et al., 2020), important inter- and intra-annual variation occurred in *E. balteatus* trap catches. Such variation can possibly be ascribed to agroclimatic conditions or biotic factors e.g., fluctuating prey abundance (Drake and Farrow, 1988; Hu et al., 2021). For aphidophagous hoverflies such as *E. balteatus*, adults deposit their eggs near aphid colonies where larvae subsequently feed until pupation; the ensuing hoverfly abundance levels are inherently shaped by aphid densities (Honěk, 1983). Aphid infestation pressure is also a prominent migration trigger for certain hoverfly species (Svensson and Janzon, 1984). Pest and natural enemy populations are thus coupled and even exhibit multi-year oscillation cycles e.g., as for the soybean aphid in North America (Rhainds et al., 2010). In our study, periodic aphid population outbreaks in mainland China likely shaped *E. balteatus* migration patterns. Hence, depending upon their voltinism, prey range and generation time, hoverfly abundance levels could provide an ‘early warning’ of pest population build-up and help target interventions in arable crops.

In migration research, an unambiguous delineation of migration pathways is complicated by insects’ flight behavior, life span and small body size. For minute insects such as hoverflies, traditional approaches such as mark-recapture methods or remote sensing have limited applicability (Hobson, 1999; Hobson and Wassenaar, 2008). Meanwhile, new methods such as trajectory analysis and endogenous markers can overcome these difficulties and advance insect migration research (Chapman et al., 2010). By coupling HYSPLIT backward trajectory analysis with stable isotope measurements, we reliably gauged *E. balteatus* migration origin and movement dynamics over hundreds of kilometers. Similar achievements have been made for other insects in the study area (Feng et al., 2009; Hu et al., 2017) and for migratory hoverflies in other parts of the globe (Raymond et al., 2013; Gao et al., 2020). Irrespective of the potential of these tracking technologies, we were unable to demarcate the exact migration routes of captured specimens or the (ephemeral, patchy) habitats that are exploited by *E. balteatus* spring migrants. Yet, considering how migrant hoverflies involve in the transport of energy, nutrients and biomass, pest regulation and pollen transfer at a macro-scale (Wotton et al., 2019), our work has major implications for environmental preservation and sustainable agri-food production in eastern Asia.

Population genomics analyses uncovered high haplotype diversity and low nucleotide diversity in *E. balteatus* - a distinguishing feature of migratory species. The lack of a pronounced *E. balteatus* genetic differentiation and genetic structure has also been recorded for other hoverfly populations (Hondelmann et al., 2005; Raymond et al., 2013) and mirrors the extent to which migration enables genetic mixing among populations from remote geographical origins. These assays also confirm the close genetic relationship between BH migrants and field-caught individuals from northeastern China. The high genetic diversity of *E. balteatus* contrasts markedly with that of crop pests or invasive species (Facon et al. 2011; Hsieh et al., 2011), possibly reflecting the species’ superior colonization ability and adaptive potential under environmental change. Elevated levels of ecological plasticity among key ecosystem service providers such as *E. balteatus* are promising in view of the recent declines in global insect diversity and abundance (Powney et al., 2019; Wotton et al., 2019; Sanchez-Bayo and Wyckhuys, 2021). Despite its important findings, our study does encounter several limitations. First, *E. balteatus* populations were primarily sampled in northern China and may not be representative of entire eastern Asia. Second, while conventional molecular markers proved to be effective for *E. balteatus* genetic analyses, next-generation sequencing (NGS) can yield a more in-depth characterization of genetic structure and evolutionary history.

With the advent of NGS approaches, molecular gut content analysis has become a powerful tool to reveal multiple trophic linkages (Pompanon et al., 2012). In our study, we validated a simple, cost-effective assay to reliably detect plant DNA in *E. balteatus* guts and to document associated (plant) feeding events. Molecular gut content analysis hereby represents advantages over other methods such as pollen analysis, to decipher insect x plant feeding relationships and to map flower visitation networks. Drawing upon this sensitive approach, we registered links to 1,012 plant species from 39 orders - far exceeding the number of host plants identified in previous studies (Lucas et al., 2018a, b). A wide spectrum of plants is thus visited by *E. balteatus* - comprising herbaceous plants, trees and several cultivated crops. Aside from confirming the role of *E. balteatus* as a key pollinator of wildflowers (Rader et al., 2020; Doyle et al., 2020), our work identified its principal (seasonal) foraging resources e.g., spring and autumn migrants relied extensively upon *Brassica L. and Artemisia L.*, respectively, these plants may be a critical resource for hoverflies, who in turn may be playing an important role in *E. balteatus* reproduction. Moreover, as exemplified in other studies, geographically-confined plant species can also help to track insect migration (e.g., Guo et al., 2022). In our study, plants endemic to central and southern China were routinely associated with spring-summer migrants. By overlaying migration dynamics with flower phenology, the underlying migration determinants can be identified e.g., individuals’ pursuit of more abundant or nutritious food resources (Chapman et al. 2011). In addition, our study reveals the extent to which migratory hoverflies mediate transcontinental pollination and may be facilitating the gene flow between geographically disjunct plant populations.

Hoverflies are prime ecosystem service providers, which not only act as the second most important pollinators after bees but equally contribute to natural biological control of a broad suite of sap-feeding crop pests. Given the countless hoverfly individuals that annually disperse within the East Asia monsoon climatic zone, the social-ecological implications of *E. balteatus* ecosystem service delivery are non-negligible i.e., contributing crop health and fruit yield (Paschke et al., 2002), while benefiting co-occurring (insect, vertebrate) pollinators, insectivores and seed feeders. Our work also constitutes a foundation for myriad follow-up experiments. For example, as several of the identified plant hosts likely mediate *E. balteatus* population build-up, further work is warranted to clarify their impacts on hoverfly nutritional ecology (Pinheiro et al., 2015) or biological control (Batuecas et al., 2021). Our findings can further aid the design of ecological engineering schemes to conserve *E. balteatus* in varying landscape contexts and to bolster its pollination and biological control services (Landis et al., 2000; Gurr et al., 2016). Lastly, NGS-based pollen identification schemes -as the ones developed in our assays- could readily be built into biodiversity monitoring programs (Reboud et al., 2022).

## Acknowledgements

We would like to express our thanks to all of the people who kindly helped us during wild samples collecting, especially Prof. Peng Wan (Institute of Plant Protection and Soil Science, Hubei Academy of Agricultural Sciences), Prof. Jian Liu (College of Agriculture, Northeast Agricultural University), Prof. Honghua Su (College of Horticulture and Plant Protection, Yangzhou University), Prof. Yutao Xiao (Agricultural Genomics Institute at Shenzhen, Chinese Academy of Agricultural Sciences), Prof. Xingya Wang (College of Plant Protection, Shenyang Agricultural University), Dr. Lili Wang (Yantai Academy of Agricultural Sciences), and Ms. Aili Li (Xinxiang Experimental Station of Chinese Academy of Agricultural Sciences). We also thank Prof. Zhiheng Wang (Peking University) for providing the atlas of plants in China. This study was funded by the Laboratory of Lingnan Modern Agriculture Project (NT2021003) and the Agricultural Science and Technology Innovation Program Cooperation and Innovation Mission (CAAS-XTCX2018022). The funders had no role in study design, data collection and analysis, decision to publish, or preparation of the manuscript.

## Author contributions

Kongming Wu conceived and designed all experiments, and led writing of the manuscript. For all experiments, Chaoxing Hu performed the backward trajectory analysis, Yongqiang Liu was responsible for morphology-based pollen grain analysis, Hui Li conducted the feeding experiments, Xianyong Zhou and Xiaokang Li assisted with development of NGS metabarcoding protocols and creation of the sequencing libraries, performed the NGS runs and assisted with data analysis, and Huiru Jia performed the rest of experiments. Huiru Jia, Xiaokang Li, Yunfei Pan and Xianyong Zhou undertook data analysis and figure design. Huiru Jia wrote the first draft of the paper, Kris A.G. Wyckhuys reviewed and edited the manuscript, and all authors made significant contributions to the final draft.

## Data availability

The sequences for population genetics studies has been deposited at GenBank, raw MiSeq data from DNA metabarcoding of gut contents has been deposited at NCBI Sequence Read Archive (SRA) (we haven’t got the accession numbers till now, so we temporarily placed all sequences in Supplementary information). All data supporting the findings of this study are available within the Article, the Extended Data and the Supplementary Information files.

## SUPPORTING INFORMATION

**Table S1.**
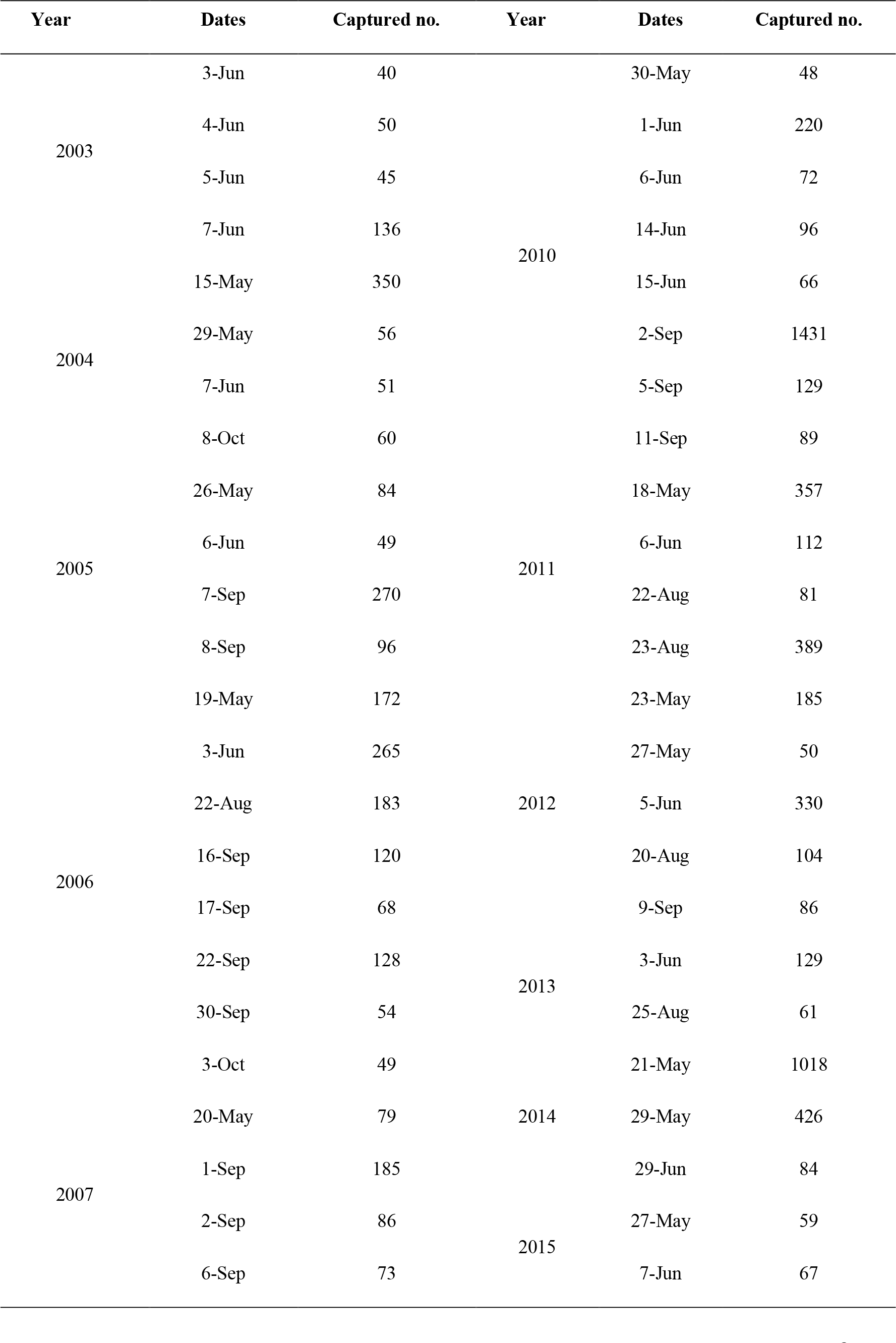
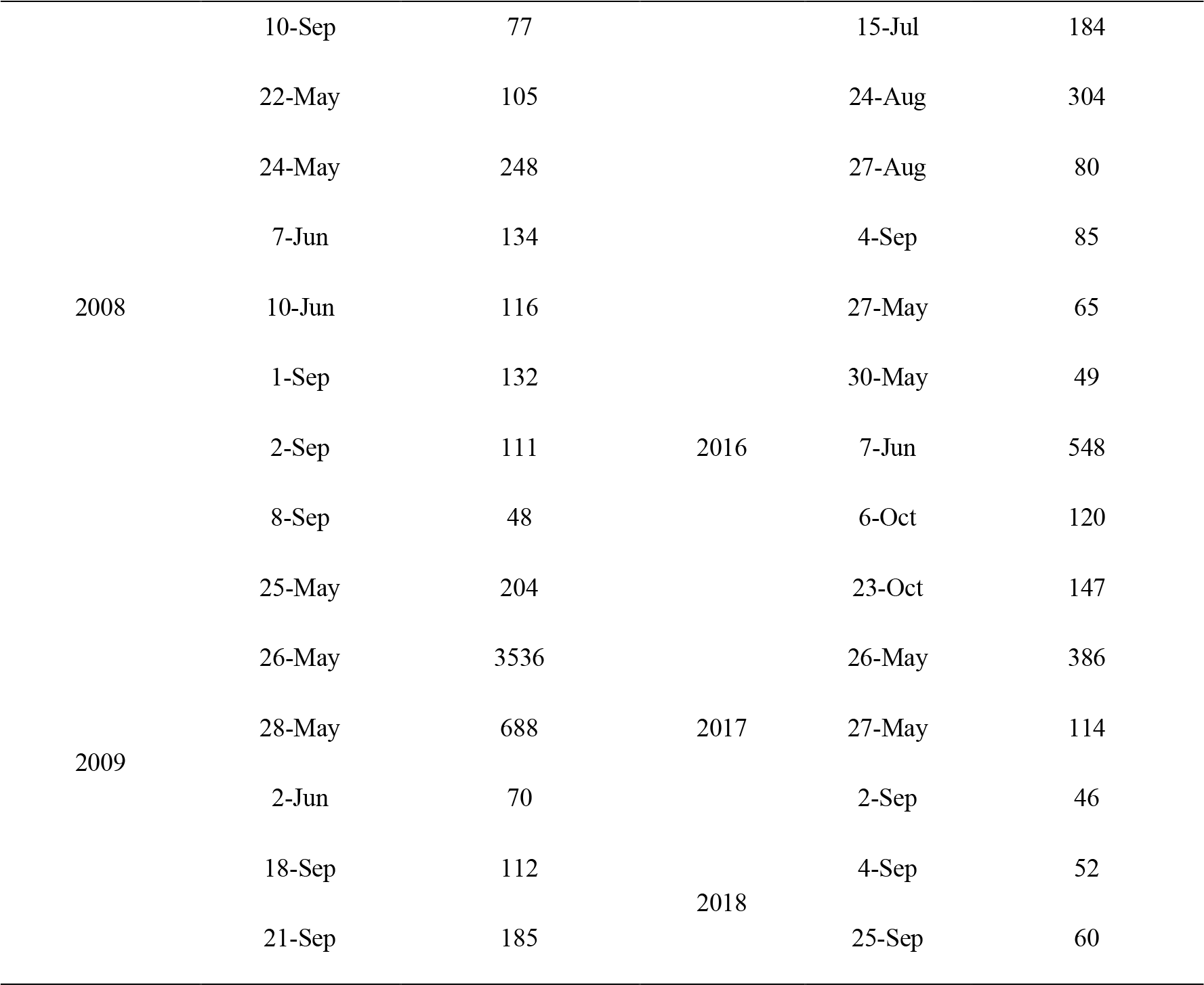
Mass migration events of E. balteatus across the Bohai Strait observed by the searchlight trapping on BH Island during 2003–2018.

**Table S2.**
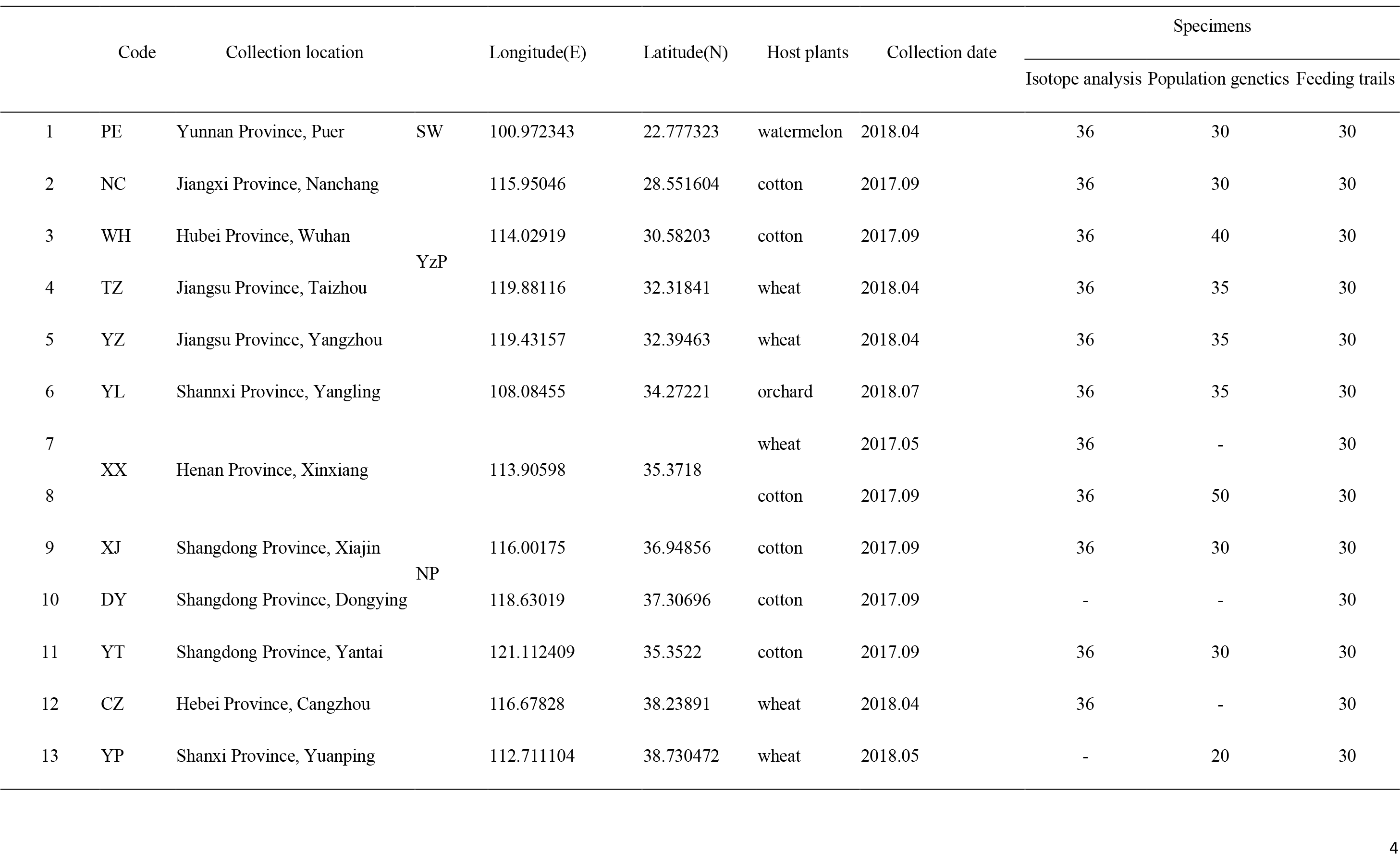
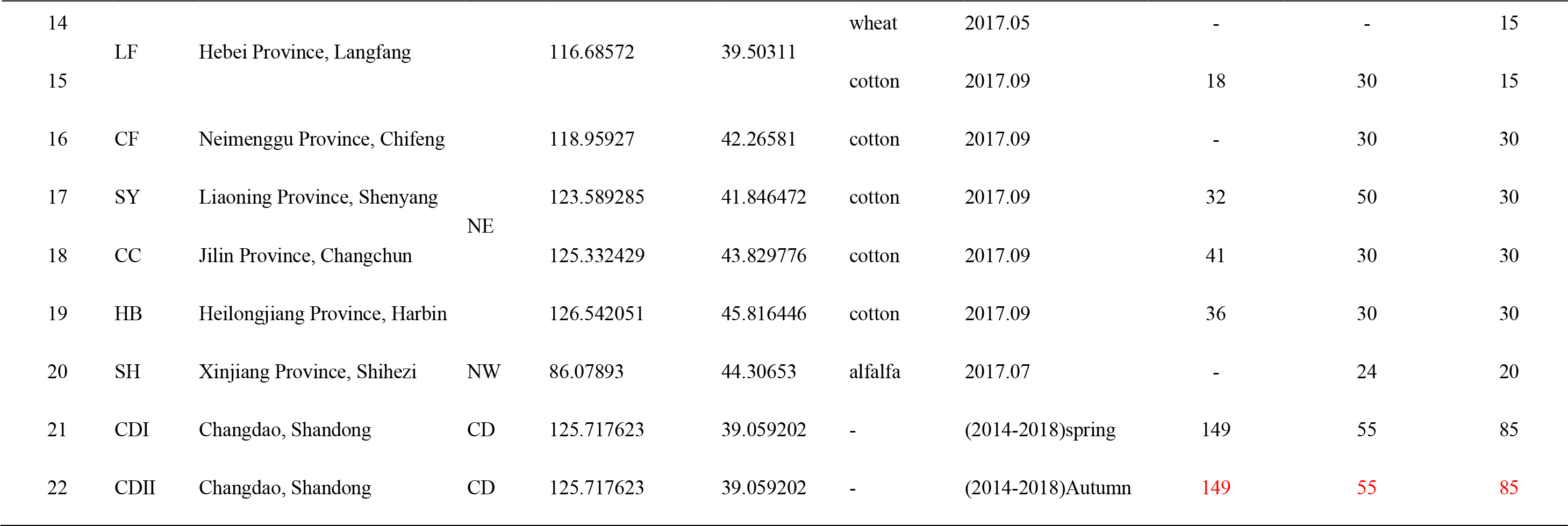
Collection information for sample in the study.

**Table S3.** Pollen carrying rate of the migratory E. balteatus hoverflies across Bohai Sea during 2014-2018.

| Sampling time |  | No. with pollen | No. adults examined | % with pollen | No. taxa |
| --- | --- | --- | --- | --- | --- |
| Spring-summer<br>migration stage | April | 7 | 42 | 17% |  |
|  | May | 101 | 367 | 28% | 28 |
|  | June | 59 | 174 | 34% |  |
|  | July | 1 | 29 | 3% | 1 |
| Autumn<br>migration stage | August | 40 | 138 | 24% |  |
|  | September | 82 | 205 | 27% | 17 |
|  | October | 30 | 59 | 50% |  |
| Total |  | 320 | 1014 | 27% | 46 |

**Table S4.**
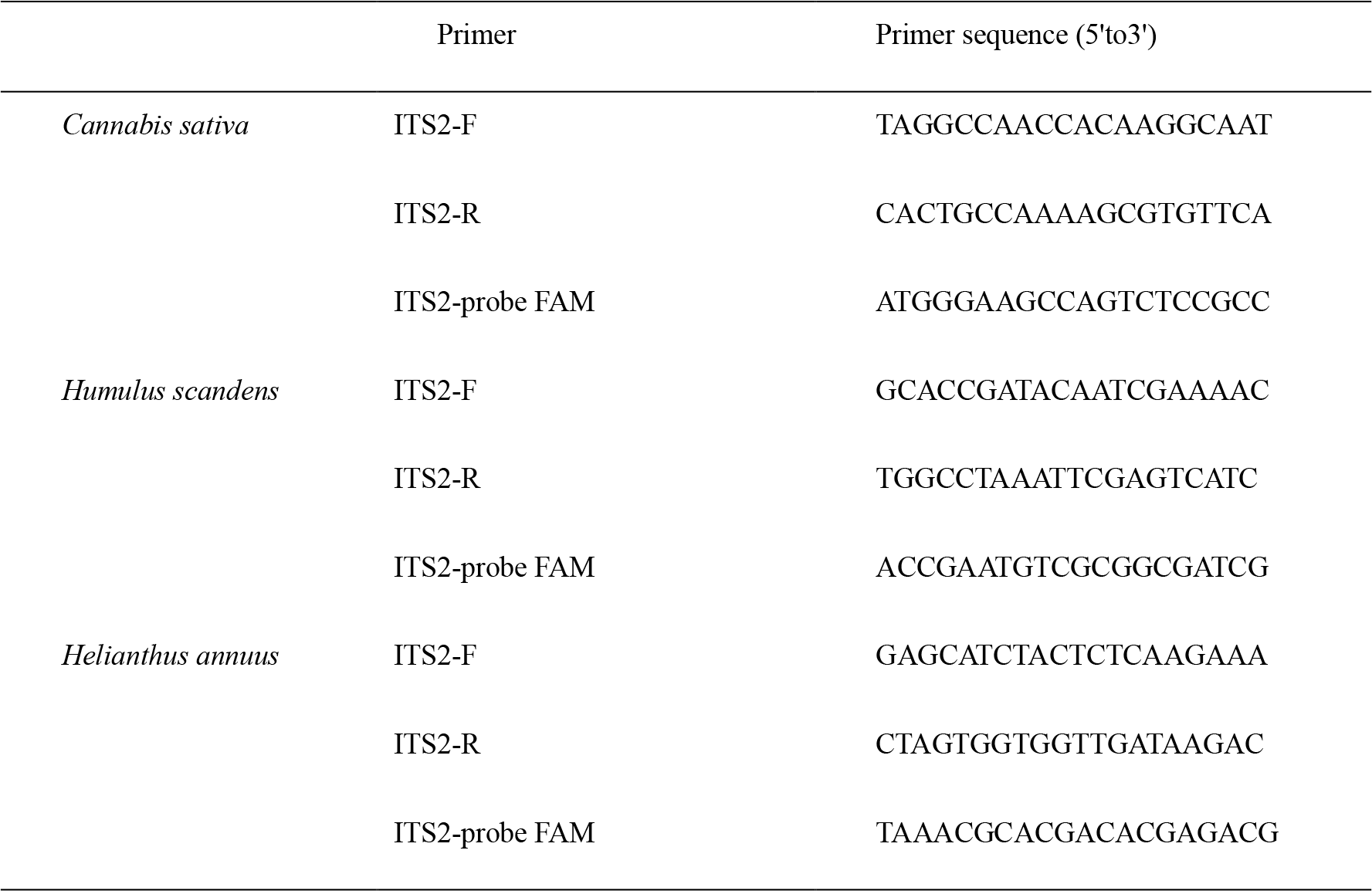
Quantitative PCR (qPCR) primers and conditions used in this study.

**Table S5.** Key parameters within E. balteatus searchlight trapping on BH Island during 2003– 2018.

| Year | First capture dd/mm<br>(no.) | Final capture dd/mm<br>(no.) | Duration of capture<br>(d) | Peak capture dd/mm<br>(no.) |
| --- | --- | --- | --- | --- |
| 2003 | 01 Jun. (10) | 21 Sep. (4) | 113 | 07 Jun. (136) |
| 2004 | 14 May (10) | 18 Oct. (8) | 158 | 15 May (350) |
| 2005 | 13 May (3) | 04 Oct. (2) | 145 | 07 Sep. (270) |
| 2006 | 17 May (16) | 20 Oct. (14) | 157 | 03 Jun. (265) |
| 2007 | 02 May (2) | 11 Oct. (1) | 163 | 01 Sep. (185) |
| 2008 | 17 May (2) | 29 Sep. (3) | 136 | 24 May (248) |
| 2009 | 05 May (2) | 29 Sep. (37) | 148 | 26 May (3536) |
| 2010 | 21 May (1) | 13 Oct. (2) | 146 | 02 Sep. (1431) |
| 2011 | 03 May (1) | 21 Sep. (11) | 142 | 23 Aug. (389) |
| 2012 | 02 May (1) | 20 Oct. (1) | 172 | 05 Jun. (330) |
| 2013 | 02 May (7) | 10 Oct. (5) | 162 | 03 Jun. (129) |
| 2014 | 01 May (13) | 28 Sep. (1) | 151 | 21 May (1018) |
| 2015 | 28 Apr.(2) | 17 Oct. (1) | 173 | 24 Aug. (304) |
| 2016 | 30 Apr.(2) | 28 Oct. (1) | 182 | 07 Jun. (548) |
| 2017 | 23 May (1) | 22 Sep. (7) | 123 | 26 May (386) |
| 2018 | 04 May (1) | 28 Sep. (10) | 148 | 25 Sep. (60) |

**Table S6.** Results of analysis of molecular variance (AMOVA) test in different populations and regions of *E.balteatus* based on Cytb and18S-28S rRNA gene.

|  | Source of variation | d.f. | Sum of squares | Variance components | Percentage of variation | Fixation Indices |
| --- | --- | --- | --- | --- | --- | --- |
| Cytb | Among populations | 17 | 16.8 | 0.012 Va | 1.84 | FST : 0.01844<br>P-value<0.02 |
|  | Within population | 512 | 327.117 | 0.6389 Vb | 98.16 |  |
|  | Total | 529 | 343.917 | 0.6509 |  |  |
| 18S-28S | Among populations | 15 | 217.7 | 0.60342 Va | 11.12 | FST : 0.11121<br>P-value<0.01 |
|  | Within population | 244 | 1176.661 | 4.82238 Vb | 88.88 |  |
|  | Total | 259 | 1394.362 | 5.42581 |  |  |

**Table S7.**
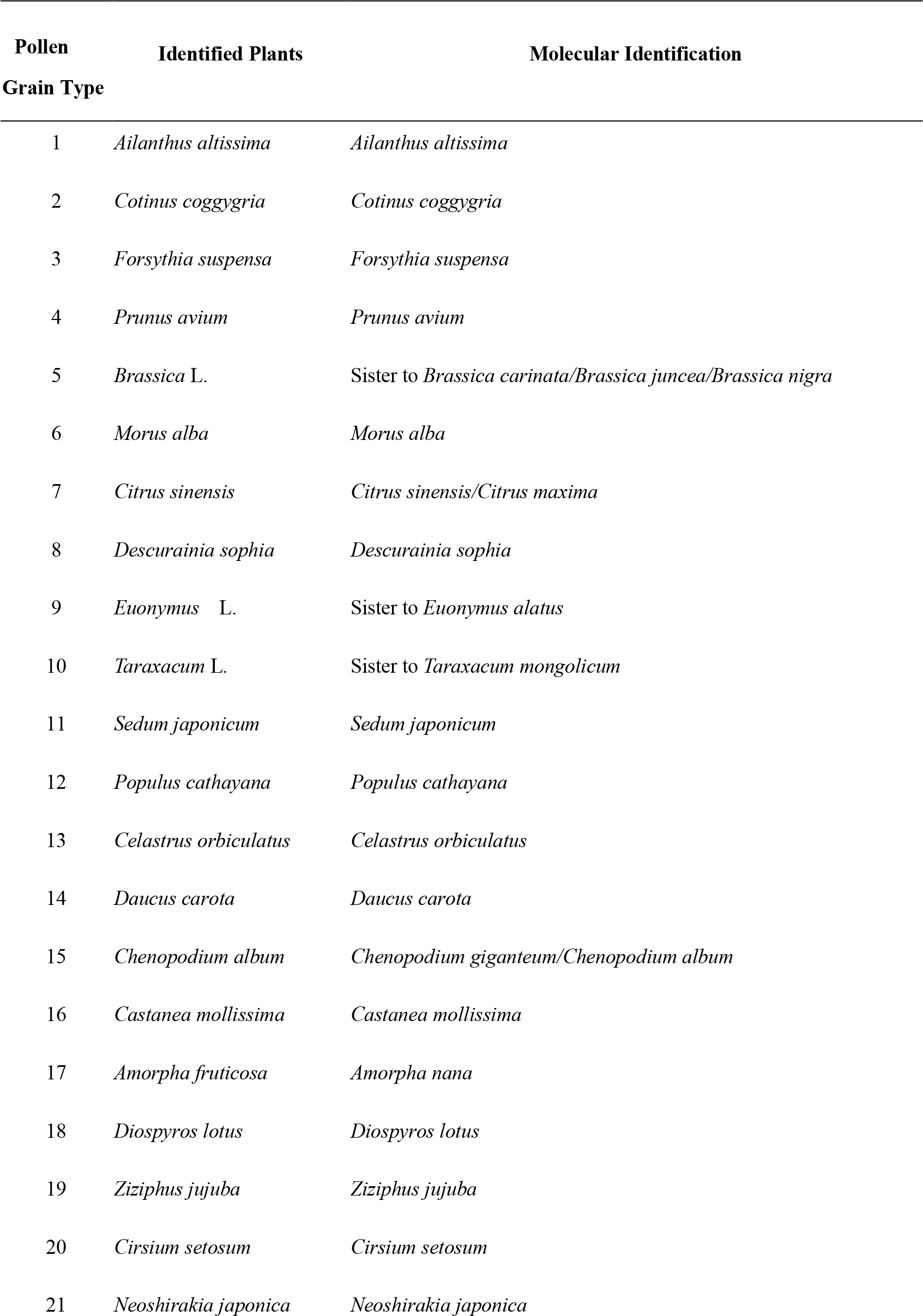
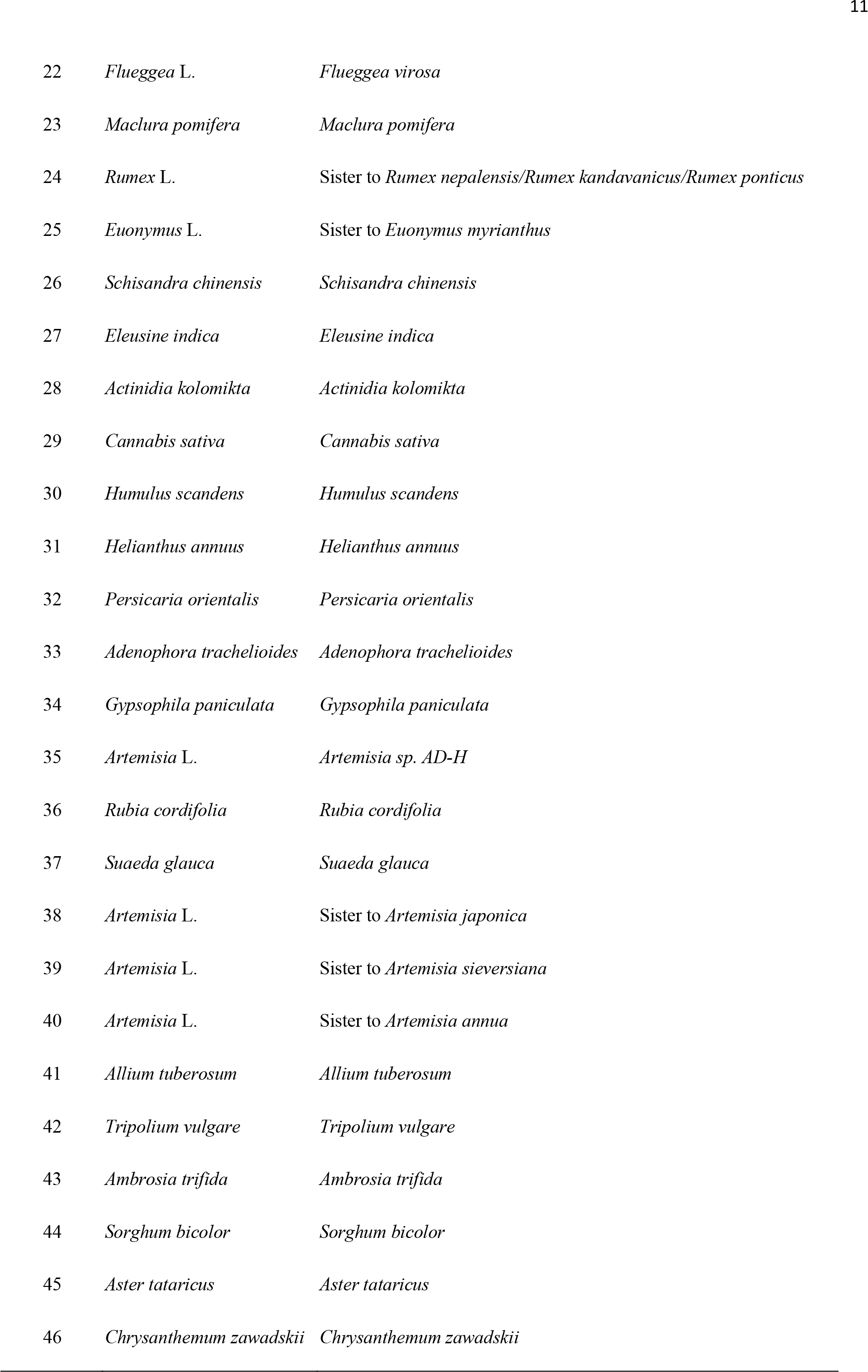
Comparative assessment of the degree of taxonomic identification obtained through either molecular or morphology-based approaches, for 46 different types of pollen grains dislocated from E. balteatus long-distance migrants collected on Beihuang Island (Bohai Sea, northeastern China). For each type of pollen grain, the highest level of taxonomic identification is indicated and contrasted between molecular and morphology-based approaches.

**Figure S1.**
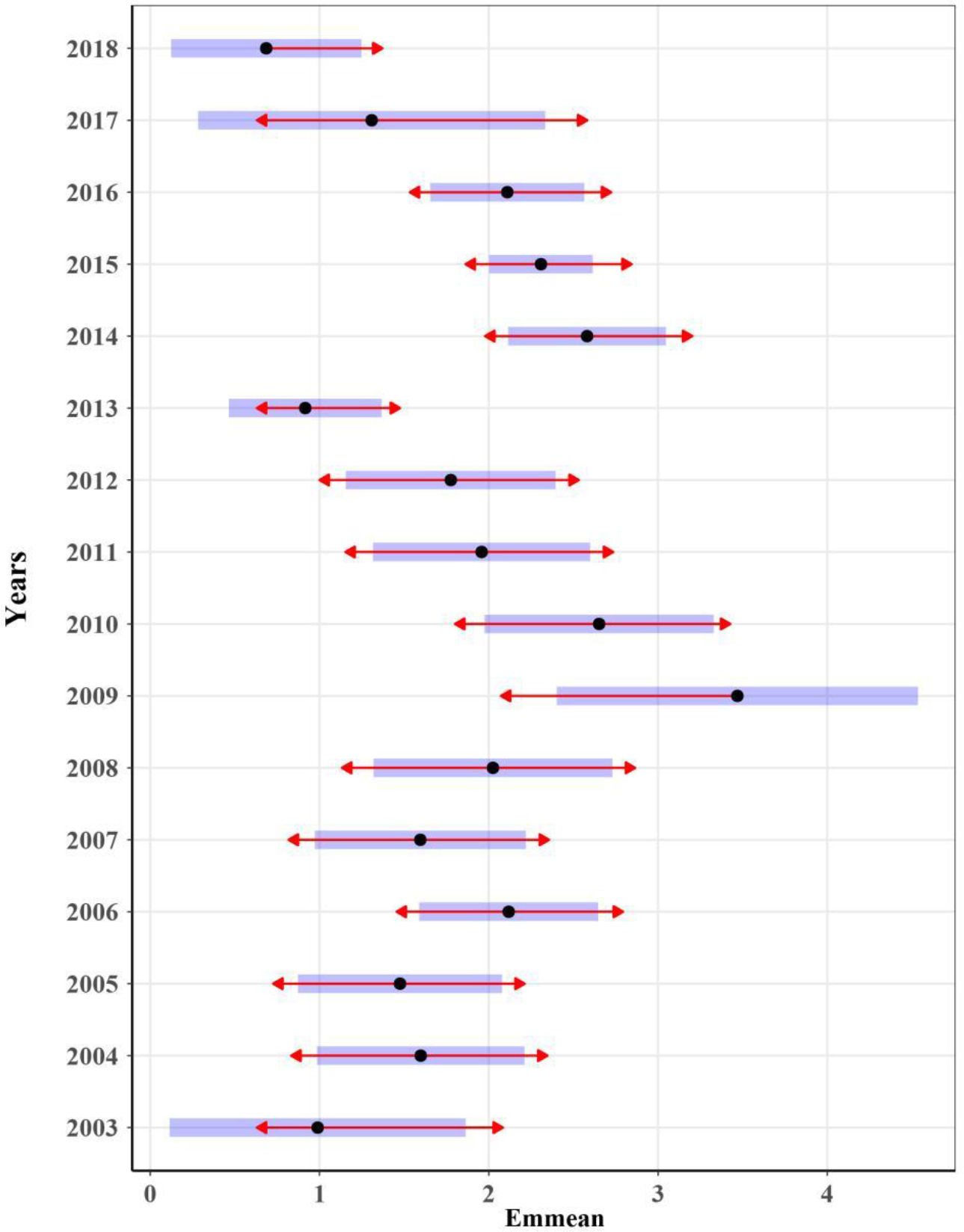
Analysis of variance on the number of *E.balteatus* captured in the searchlight trap on BH Island from May to October 2003–2018 by comparing EMMs. The blue bars are confidence intervals for the EMMs, and the red arrows are for the comparisons among them. If an arrow from one mean overlaps an arrow from another group, the difference is not “significant,” based on the adjust setting (which defaults to “tukey”) and the value of alpha (which defaults to 0.05).

**Figure S2.**
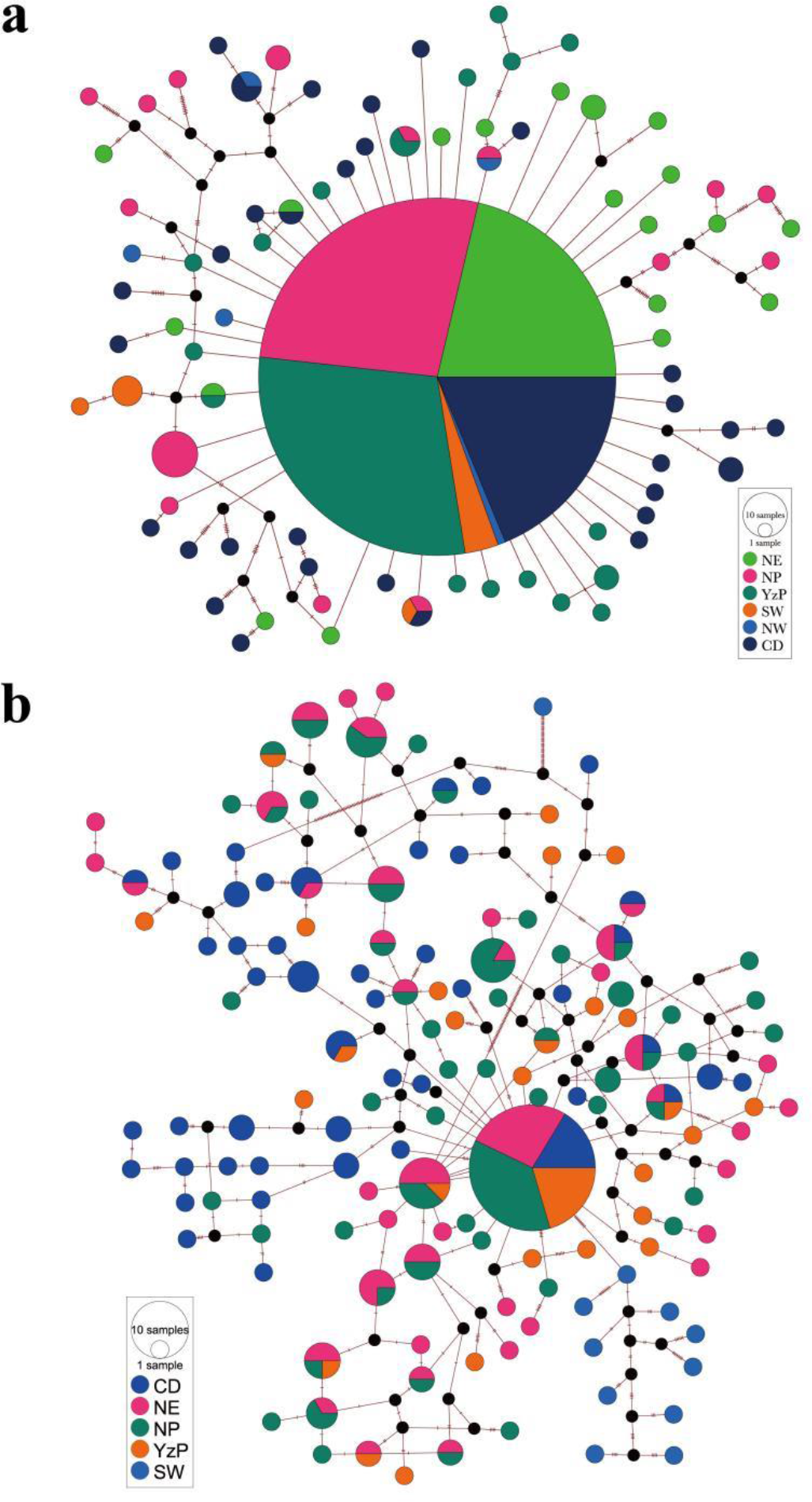
Median-joining haplotype network of the gene *CytB* (a) and 18S-28S rRNA gene (b). Circle areas indicated the proportion of haplotype frequencies, while colored portions represent the proportions of the same haplotype that occurs in each region.

**Figure S3.**
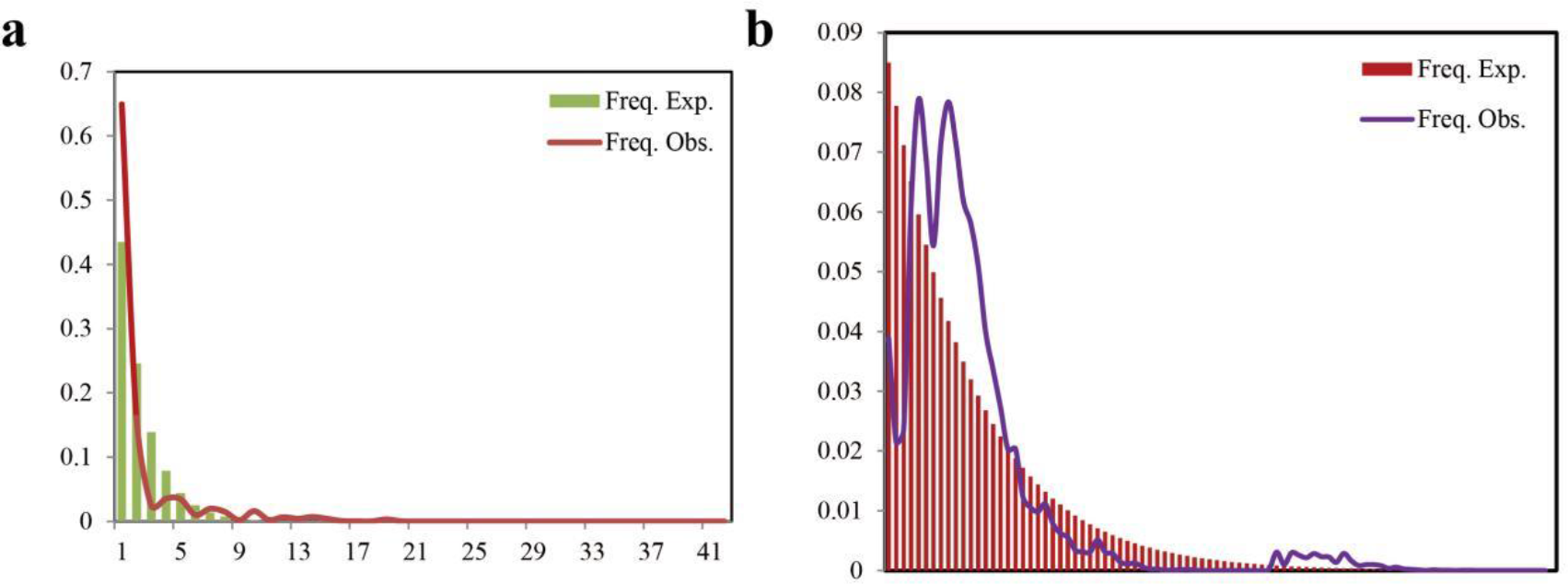
Mismatch distribution of the Cytb and18S-28S rRNA gene in *E.balteatus* populations.

**Figure S4.**
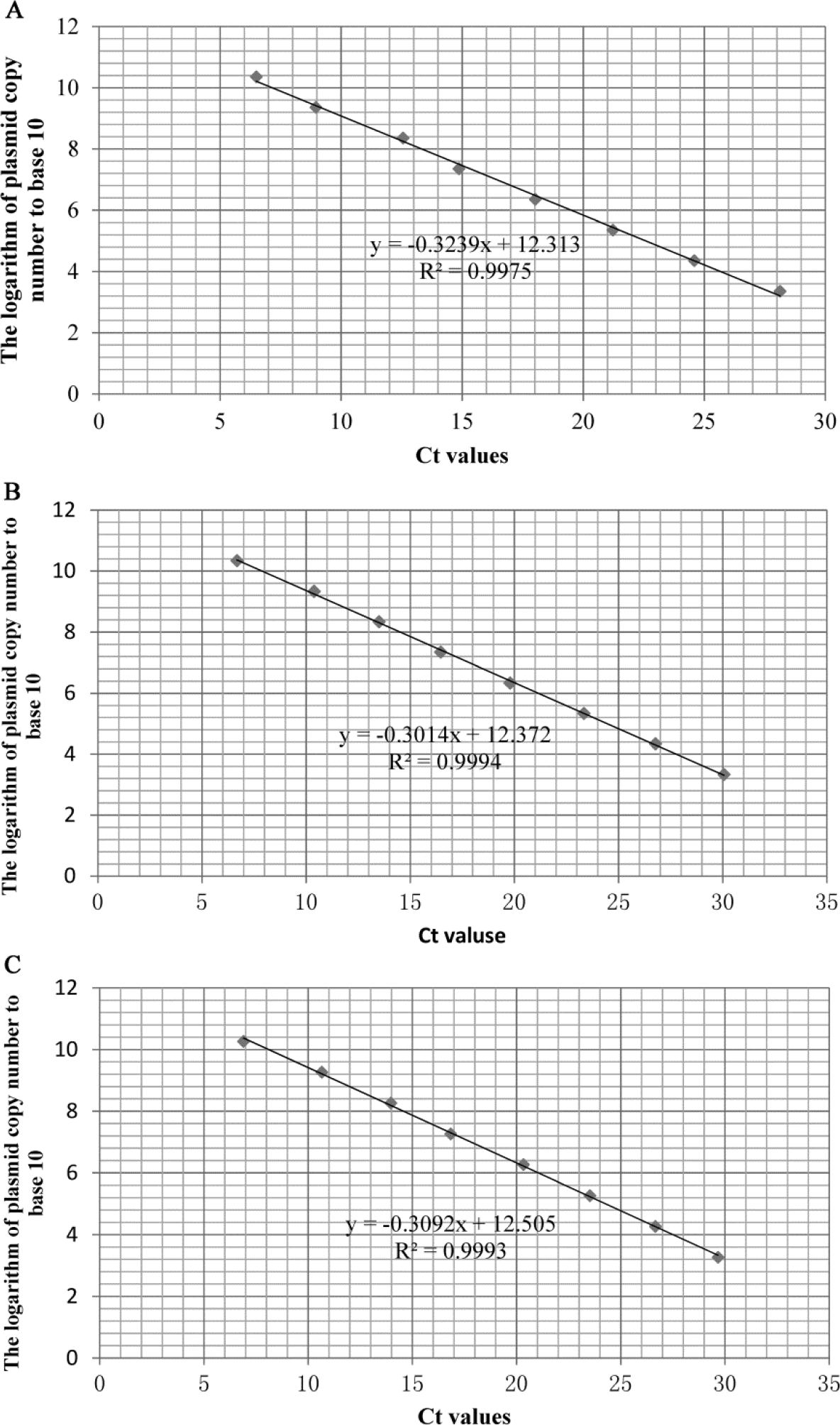
The respective standard curve equations of the three plants in the feeding experiments. A:*Cannabis sativa*, B:*Humulus scandens* C: *Helianthus annuus*

**Figure S5.**
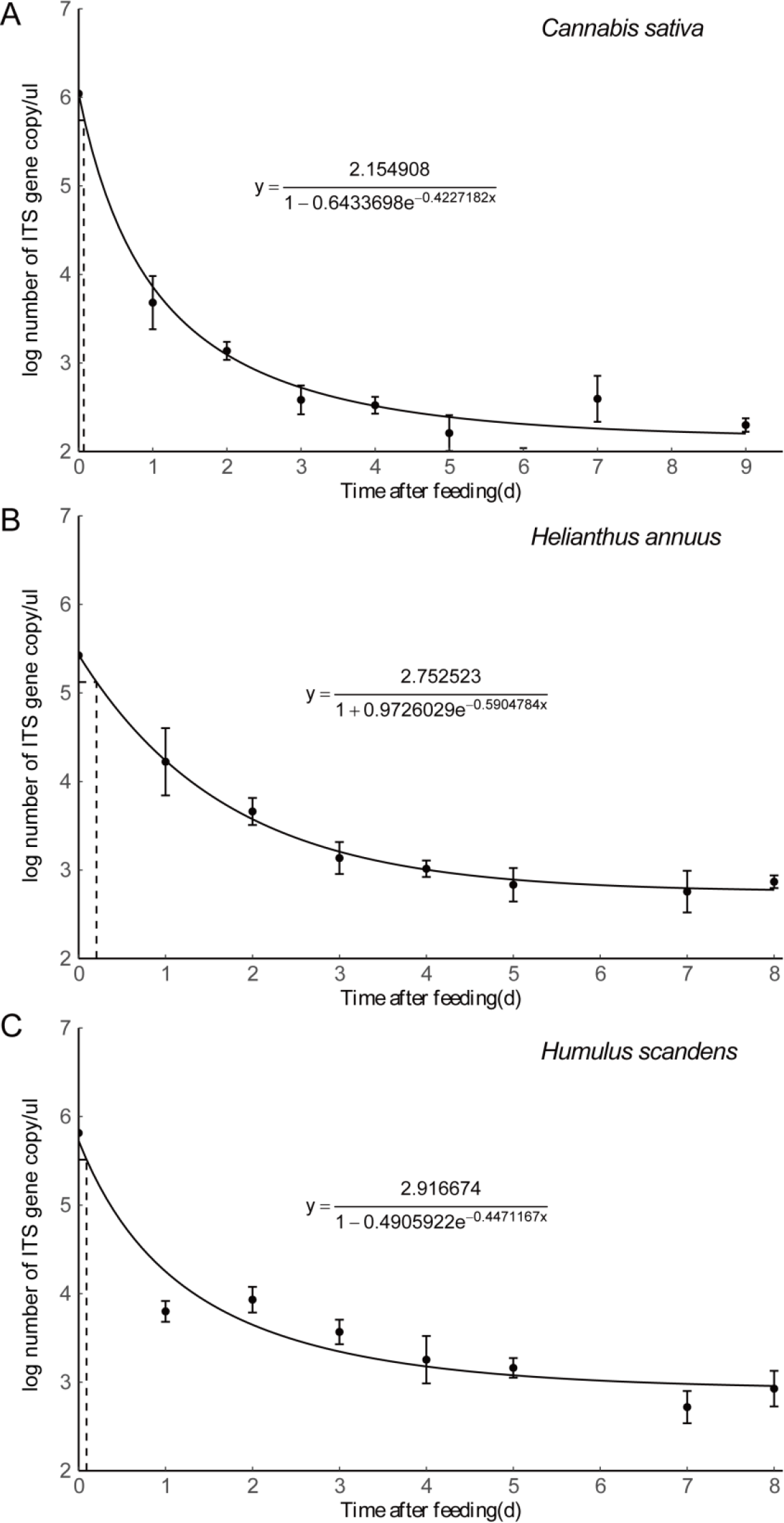
Detection of three different plant DNA in the guts of *E.balteatus* adults at different times after ingestion by qPCR analysis. Error bars at each point on the curves represent the standard error of replicates.

## References

1. Anderson RC. 2009. Do dragonflies migrate across the western Indian Ocean?. Journal of tropical Ecology 25:347–358.

2. Aubert J, Goeldlin de Tiefenau P. 1981. Observations sur les migrations de Syrphides (Dipt.) dans les Alpes de Suisse occidentale. Mitteilungen der Schweizerischen Entomologischen Gesellschaft 54: 377–388.

3. Bandelt HJ, Forster P, Röhl A. 1999. Median-joining networks for inferring intraspecific phylogenies. Molecular biology and evolution 16:37–48.

4. Batuecas I, Agustí N, Castañé C, Alomar O. 2021. Molecular tracking of insect dispersal to verify arthropod predator movement from an alfalfa field to a peach orchard. Biological Control 158:104506.

5. Beerli P. 2006. Comparison of Bayesian and maximum-likelihood inference of population genetic parameters. Bioinformatics 22:341–345.

6. Branquart E, Hemptinn J. 2000. Selectivity in the exploitation of floral resources by hoverflies (Diptera: Syrphinae). Ecography 23:732–742.

7. Brooks ME, Kristensen K, Van Benthem KJ, Magnusson A, Berg CW, Nielsen A, Skaug HJ, Machler M, Bolker, BM. 2017. glmmTMB balances speed and flexibility among packages for zero-inflated generalized linear mixed modeling. The R journal 9:378–400.

8. Camacho C, Coulouris G, Avagyan V, Ma N, Papadopoulos J, Bealer K, Madden TL. 2009. BLAST+: architecture and applications. BMC bioinformatics 10:1–9.

9. Chapman JW, Bell JR, Burgin LE, Reynolds DR, Pettersson LB, Hill JK, Bonsall MB, Thomas JA. 2012. Seasonal migration to high latitudes results in major reproductive benefits in an insect. PNAS 109:14924–14929 .

10. Chapman JW, Drake VA, Reynolds DR. 2011. Recent insights from radar studies of insect flight. Annual review of entomology 56: 337–356.

11. Chapman JW, Nesbit RL, Burgin LE, Reynolds DR, Smith AD, Middleton DR, Hill JK. 2010. Flight Orientation Behaviors Promote Optimal Migration Trajectories in High-Flying Insects. Science 327:682–685.

12. Chapman JW, Reynolds DR, Wilson K. 2015. Review and long-range seasonal migration in insects: mechanisms, evolutionary drivers and ecological consequences. Ecology letters 18:287–302.

13. Chen PH, Pan YB, Chen RK. 2008. High-throughput procedure for single pollen grain collection and polymerase chain reaction in plants. Journal of Integrative Plant Biology 50:375–383.

14. Cheng T, Xu C, Lei L, Li C, Zhang Y, Zhou S. 2016. Barcoding the kingdom Plantae: new PCR primers for ITS regions of plants with improved universality and specificity. Molecular Ecology Resources 16:138–149.

15. Dällenbach LJ, Glauser A. Lim KS, Chapman JW, Menz MH. 2018. Higher flight activity in the offspring of migrants compared to residents in a migratory insect. Proceedings of the Royal Society B 285:20172829.

16. Doyle T, Hawkes WL, Massy R, Powney GD, Menz MH, Wotton KR. 2020. Pollination by hoverflies in the Anthropocene. Proceedings of the Royal Society B 287:20200508.

17. Drake VA, Farrow RA. 1988. The influence of atmospheric structure and motions on insect migration. Annual review of entomology 33:183–210.

18. Drake, V A, Farrow RA. 1988. The influence of atmospheric structure and motions on insect migration. Annual review of entomology 33:183–210.

19. Excoffier L, Lischer HE. 2010. Arlequin suite ver 3.5: a new series of programs to perform population genetics analyses under Linux and Windows. Molecular Ecology Resources 10:564–567.

20. Facon B, Hufbauer RA, Tayeh A, Loiseau A, Lombaert E, Vitalis R, Thomas G, Lundgren JG, Estoup A. 2011. Inbreeding depression is purged in the invasive insect *Harmonia axyridis*. Current Biology 21: 424–427.

21. Fay MF, Swensen SM, Chase MW. 1997. Taxonomic affinities of *Medusagyne oppositifolia* (Medusagynaceae). Kew Bulletin 52:111–120.

22. Fazekas AJ, Burgess KS, Kesanakurti PR, Graham SW, Newmaster SG, Husband BC, Percy DM, Hajibabaei M, Barrett SC. 2008. Multiple multilocus DNA barcodes from the plastid genome discriminate plant species equally well. PLoS ONE 3:e2802.

23. Feng H, Wu X, Wu B, Wu K. 2009. Seasonal migration of *Helicoverpa armigera* (Lepidoptera: Noctuidae) over the Bohai Sea. Journal of Economic Entomology 102:95–104.

24. Feng HQ, Wu KM, Cheng DF, Guo YY. 2003. Radar observations of the autumn migration of the beet armyworm *Spodoptera exigua* (Lepidoptera: Noctuidae) and other moths in northern China. Bulletin of Entomological Research 93:115– 124 .

25. Feng HQ, Wu KM, Ni YX, Cheng DF, Guo YY. 2006. Nocturnal migration of dragonflies over the Bohai Sea in northern China. Ecological Entomology 31: 511–520 .

26. Finch JT, Cook JM. 2020. Flies on vacation: evidence for the migration of Australian syrphidae (diptera). Ecological Entomology 45:896–900.

27. Fu YX. 1997. Statistical tests of neutrality of mutations against population growth, hitchhiking and background selection. Genetics 147:915–925.

28. Gao B, Wotton KR, Hawkes WL, Menz MH, Reynolds DR, Zhai BP, Gao H, Chapman JW. 2020. Adaptive strategies of high-flying migratory hoverflies in response to wind currents. Proceedings of the Royal Society B 287:20200406.

29. García-Robledo C, Erickson DL, Staines CL, Erwin TL, Kress WJ. 2013. Tropical plant-herbivore networks: Reconstructing species interactions using DNA barcodes. PLoS ONE 8:e52967.

30. Gariepy TD, Kuhlmann U, Gillott C, Erlandson M. 2007. Parasitoids, predators and PCR: the use of diagnostic molecular markers in biological control of Arthropods. Journal of Applied Entomology 131:225–240.

31. Guo J, Fu X, Zhao S, Shen X, Wyckhuys KA, Wu K. 2020. Long-term shifts in abundance of (migratory) crop-feeding and beneficial insect species in northeastern Asia. Journal of Pest Science 93:583–594.

32. Guo J, Liu Y, Jia H, Chang H, Wu K. 2022. Visiting Plants of *Mamestra brassicae* (Lepidoptera: Noctuidae) Inferred From Identification of Adhering Pollen Grains. Environmental Entomology. DOI: https://doi.org/10.1093/ee/nvab145.

33. Gurr GM, Lu Z, Zheng X, Xu H, Zhu P, Chen G, Yao X, Cheng J, Zhu Z, Catindig JL, Villareal S. 2016. Multi-country evidence that crop diversification promotes ecological intensification of agriculture. Nature Plants 2:1–4.

34. Han J, Zhu Y, Chen X, Liao B, Yao H, Song J, Chen S, Meng F. 2013. The short ITS2 sequence serves as an efficient taxonomic sequence tag in comparison with the full-length ITS. BioMed research international 2013:741476.

35. Hawkins J, de Vere N, Griffith A, Ford CR. 2015. Using DNA Metabarcoding to Identify the Floral Composition of Honey: A New Tool for Investigating Honey Bee Foraging Preferences. PLoS ONE 10: e0134735.

36. Hobson KA, Wassenaar LI. 2008. Tracking animal migration using stable isotopes. Handbook of Terrestrial Ecology Series. Academic Press, London, UK.

37. Hondelmann P, Borgemeister C, Poehling HM. 2005. Restriction fragment length polymorphisms of different DNA regions as genetic markers in the hoverfly *Episyrphus balteatus* (Diptera: Syrphidae). Bulletin of entomological research 95:349–359.

38. Honěk A. 1983. Factors affecting the distribution of larvae of aphid predators (Col., Coccinellidae and Dipt., Syrphidae) in cereal stands. Zeitschrift f ü r angewandte Entomologie 95:336–345.

39. Hsieh CH, Chiang YH, Ko CC. 2011. Population genetic structure of the newly invasive Q biotype of *Bemisia tabaci* in Taiwan. Entomologia experimentalis et applicata 138:263–271.

40. Hu CX, Fu XW, Wu KM. 2017. Seasonal migration of white-backed planthopper *Sogatella furcifera* Horváth (Hemiptera: Delphacidae) over the Bohai Sea in northern China. Journal of Asia-Pacific Entomology 20:1358–1363.

41. Hu G, Lim KS, Horvitz N, Clark SJ, Reynolds DR, Sapir N, Chapman JW. 2016. Mass seasonal bioflows of high-flying insect migrants. Science 354: 1584–1587.

42. Hu G, Stefanescu C, Oliver T H, Roy DB, Brereton T, Van Swaay C, Reynolds DR, Chapman JW. 2021. Environmental drivers of annual population fluctuations in a trans-Saharan insect migrant. PNAS 118: e2102762118.

43. Huestis DL, Dao A, Diallo M, Sanogo ZL, Samake D, Yaro AS, et al. 2019. Windborne long-distance migration of malaria mosquitoes in the Sahel. Nature 574:404–408.

44. Jones GD, Jones, SD. 2001. The uses of pollen and its implication for entomology. Neotropical Entomology 30: 341–350.

45. Kim KS, Sappington TW. 2013. Population genetics strategies to characterize long-distance dispersal of insects. Journal of Asia-Pacific Entomology 16:87–97.

46. Lack D, Lack E. 1951. Migration of insects and birds through a Pyrenean pass.The Journal of Animal Ecology 20:63–67.

47. Landis DA, Wratten SD, Gurr GM. 2000. Habitat management to conserve natural enemies of arthropod pests in agriculture. Annual review of entomology 45: 175–201.

48. Li H, Jiang SS, Zhang HW, Geng T, Wyckhuys KA, Wu KM. 2021. Two-way predation between immature stages of the hoverfly *Eupeodes corollae* and the invasive fall armyworm (*Spodoptera frugiperda* JE Smith). Journal of Integrative Agriculture 20:829–839.

49. Li TQ, Cao HJ, Kang MS, Zhang ZX, Zhao N, Zhang H. 2010. Pollen Flora of China, Woody Plants by SEM. Science Press: Beijing, China.

50. Li W, Wu M, Lian, Z. 2009. Progress in research on the Syrphidae in China. Chinese Journal of Applied Entomology 46:861–865.

51. Liu Y, Fu X, Mao L, Xing Z, Wu K. 2016. Host plants identification for adult Agrotis ipsilon, a long-distance migratory insect. International journal of molecular sciences 17:851–863.

52. Lucas A, Bodger O, Brosi BJ, Ford CR, Forman DW, Greig C, Hegarty M, Jones L, Penelope JN, de Vere Natasha. 2018a. Floral resource partitioning by individuals within generalised hoverfly pollination networks revealed by DNA metabarcoding. Scientific reports 8:5133.

53. Lucas A, Bodger O, Brosi BJ, Ford CR., Forman DW, Greig C, Hegarty M, Neyland PJ, de Vere Natasha. 2018b. Generalisation and specialisation in hoverfly (Syrphidae) grassland pollen transport networks revealed by DNA metabarcoding. Journal of Animal Ecology 87:1008–1021.

54. Ma DW, Zhang CH, Gao SZ, Ma N, Liu, HH, Zhang YP, Sun L. 1999. Pollen Flora of China Vegetables by SEM. China Agriculture Press.

55. Mengual X, Ståhls G, Rojo S. 2015. Phylogenetic relationships and taxonomic ranking of pipizine flower flies (Diptera: Syrphidae) with implications for the evolution of aphidophagy. Cladistics 31:491–508.

56. Menz MHM, Brown BV, Wotton KR. 2019. Quantification of migrant hoverfly movements (Diptera: Syrphidae) on the West Coast of North America. Royal Society open science 6:190153.

57. Michalik B, Brust V, Hüppop O. 2020. Are movements of daytime and nighttime passerine migrants as different as day and night?. Ecology and Evolution 10: 11031–11042.

58. Pang X, Shi L, Song J, Chen X, Chen S. 2013. Use of the potential DNA barcode ITS2 to identify herbal materials. Journal of natural medicines 67:571–575.

59. Paschke M, Abs C, Schmid B. 2002. Effects of population size and pollen diversity on reproductive success and offspring size in the narrow endemic *Cochlearia bavarica* (Brassicaceae). American Journal of Botany 89:1250–1259.

60. Pinheiro LA, Torres LM, Raimundo J, Santos SA. 2015. Effects of pollen, sugars and honeydew on lifespan and nutrient levels of *Episyrphus balteatus*. BioControl 60:47–57.

61. Pompanon F, Deagle, BE, Symondson WO, Brown DS, Jarman SN, Taberlet P. 2012. Who is eating what: diet assessment using Next Generation Sequencing. Molecular Ecology 21:1931–1950.

62. Powney GD, Carvell C, Edwards M, Morris RK, Roy HE, Woodcock BA, Isaac NJ. 2019. Widespread losses of pollinating insects in Britain. Nature communications 10:1018.

63. Prosser SW, Hebert PD. 2017. Rapid identification of the botanical and entomological sources of honey using DNA metabarcoding. Food Chemistry 214:183–191.

64. R Core Team. 2020. R: A Language and Environment for Statistical Computing(manual). R Foundation for Statistical Computing,Vienna, Austria.

65. Rader R, Cunningham SA, Howlett, BG, Inouye DW. 2020. Non-bee insects as visitors and pollinators of crops: Biology, ecology, and management. Annual review of entomology 65:391–407.

66. Raymond L, Plantegenest M, Vialatte A. 2013. Migration and dispersal may drive to high genetic variation and significant genetic mixing: the case of two agriculturally important, continental hoverflies (*Episyrphus balteatus* and *Sphaerophoria scripta*). Molecular Ecology 22:5329–5339.

67. Raymond L, Vialatte A, Plantegenest M. 2014. Combination of morphometric and isotopic tools for studying spring migration dynamics in *Episyrphus balteatus*. Ecosphere 5:1–16.

68. Reboud X, Poggi S, Bohan DA. 2022. Effective biodiversity monitoring could be facilitated by networks of simple sensors and a shift to incentivising results. In Advances in Ecological Research. Academic Press.

69. Rhainds M, Yoo HJS, Kindlmann P, Voegtlin D, Castillo D, Rutledge C, Sadof C, Yaninek S, O’Neil RJ. 2010. Two-year oscillation cycle in abundance of soybean aphid in Indiana. Agricultural and Forest Entomology 12:251–257.

70. Rogers AR, Harpending H. 1992. Population growth makes waves in the distribution of pairwise genetic differences. Molecular biology and evolution 9: 552–569.

71. Rozas J, Ferrer-Mata A, Sánchez-DelBarrio JC, Guirao-Rico S, Librado P, Ramos-Onsins SE, Sánchez-Gracia A. 2017. DnaSP 6: DNA sequence polymorphism analysis of large data sets. Molecular biology and evolution 34: 3299–3302 .

72. Sánchez-Bayo F, Wyckhuys KA. 2021. Further evidence for a global decline of the entomofauna. Austral Entomology 60:9–26.

73. Satterfield DA, Sillett TS, Chapman JW, Altizer S, Marra PP. 2020. Seasonal insect migrations: Massive, influential, and overlooked. Frontiers in Ecology and the Environment 18: 335–344.

74. Searle SR, Speed FM, Milliken GA. 1980. Population Marginal Means in the Linear Model: An Alternative to Least Squares Means. The American Statistician, 34:216–221.

75. Simmons RB, Weller SJ. 2001. Utility and evolution of *cytochrome b* in insects. Molecular phylogenetics and evolution 20:196–210.

76. Staudacher K, Wallinger C, Schallhart N, Traugott M. 2011. Detecting ingested plant DNA in soil-living insect larvae. Soil Biology and Biochemistry, 43:346–350.

77. Stefanescu C, Alarcón M, Àvila A. 2007. Migration of the painted lady butterfly, *Vanessa cardui*, to north-eastern Spain is aided by African wind currents. Journal of Animal Ecology, 76, 888–898.

78. Stein AF, Draxler RR, Rolph, GD, Stunder BJ, Cohen, M.D, Ngan F. 2015. NOAA’s HYSPLIT atmospheric transport and dispersion modeling system. Bulletin of the American Meteorological Society 96:2059–2077.

79. Sun XX, Hu CX, Jia HR, Wu QL, Shen XJ, Zhao SY, Jiang YY, Wu KM. 2021. Case study on the first immigration of fall armyworm, Spodoptera frugiperda invading into China. Journal of Integrative Agriculture 20:664–672.

80. Svensson BG, Janzon LA. 1984. Why does the hoverfly *Metasyrphus corollae* migrate?. Ecological entomology 9:329–335.

81. Tajima F. 1989. Statistical method for testing the neutral mutation hypothesis by DNA polymorphism. Genetics 123:585–595.

82. Tamura K, Stecher G, Peterson D, Filipski, A, Kumar, S. 2013. MEGA6: molecular evolutionary genetics analysis version 6.0. Molecular biology and evolution 30:2725–2729.

83. Tenhumberg B, Poehling HM. 1995. Syrphids as natural enemies of cereal aphids in Germany: aspects of their biology and efficacy in different years and regions. Agriculture, Ecosystems & Environment 52:39–43.

84. Tenhumberg, B. 1995. Estimating predatory efficiency of *Episyrphus balteatus* (Diptera: Syrphidae) in cereal fields. Environmental Entomology 24:687–691.

85. Wallinger C, Staudacher K, Schallhart N, Peter E, Dresch P, Juen A, Traugott M. 2013. The effect of plant identity and the level of plant decay on molecular gut content analysis in a herbivorous soil insect. Molecular Ecology Resources 13:75–83.

86. Wang YQ. 2014. MeteoInfo: GIS software for meteorological data visualization and analysis. Meteorological Applications 21:360–368 .

87. Wassenaar LI, Hobson KA. 1998. Natal origins of migratory monarch butterflies at wintering colonies in Mexico: new isotope evidence. PNAS 95:15436–15439.

88. Wotton KR, Gao B, Menz MH, Morris RK, Ball SG, Lim KS. Don R. Reynolds, Hu G, Chapman JW. 2019. Mass seasonal migrations of hoverflies provides extensive pollination and crop protection services. Current Biology 29:2167–2173.

89. Zeng J, Liu Y, Zhang H, Liu J, Jiang Y, Wyckhuys KA, Wu K. 2020. Global warming modifies long-distance migration of an agricultural insect pest. Journal of Pest Science 9:569–581.

